# CryoGO enables high-resolution structural profiling of endogenous cellular macromolecules

**DOI:** 10.64898/2026.06.26.734647

**Authors:** Yujie Li, Yuekang Zhang, Chunxiang Wu, Chenghao Zhu, Yong Xiong

**Affiliations:** Department of Molecular Biophysics and Biochemistry, Yale University, New Haven, CT, USA

## Abstract

Resolving macromolecular structures within their native cellular environment is essential for connecting molecular architecture to physiological function, yet the field lacks accessible, high-throughput methods for generating suitable specimens from cells. Here, we introduce cryoGO (on-Grid Opening cryo-electron microscopy), a simple, rapid, and scalable strategy that mechanically opens cells directly on EM grids to produce cell-derived specimens for high-resolution single-particle cryo-EM. This approach enables near-atomic structure determination of diverse endogenous macromolecular assemblies spanning a broad molecular-weight range. We show that cryoGO captures the rich ribosomal conformational and compositional landscape while preserving physiological state distributions and aspects of spatial heterogeneity originating from cells. Furthermore, this rapid workflow enables time-resolved structural profiling, capturing both long-term cellular adaptation and acute remodeling on timescales of seconds. Compatible with standard cryo-EM infrastructure and diverse biological samples, while requiring only small numbers of cells, cryoGO lowers the technical barrier to high-resolution native structural biology.

## Introduction

Single-particle cryo-electron microscopy (cryo-EM) has transformed structural biology by enabling near-atomic-resolution visualization of purified macromolecular complexes^1–3^. However, many cellular machines function within crowded and spatially organized environments, where local molecular composition and physiological conditions shape their structures, interactions, and conformational states^4,5^. Native structural biology approaches that preserve cellular context therefore provide an important opportunity to bridge high-resolution structural biology with cell biology^6,7^. Cryo-electron tomography (cryo-ET) has been the dominant approach for visualizing macromolecular assemblies within cells, but its throughput and achievable resolution often limit routine high-resolution structural determination^8–11^. On the other hand, *in situ* single-particle cryo-EM provides a complementary strategy by applying tilt-free data collection directly to native cellular samples, allowing high-resolution reconstructions of endogenous complexes^12–15^.

Despite this promise, native cryo-EM remains technically difficult because intact cells are generally too thick for efficient electron transmission. Cryo-focused ion beam (cryo-FIB) milling can physically thin vitrified cells into lamellae suitable for imaging while preserving the native cellular environment with high fidelity^16–18^. However, its broader application is limited by low throughput, specialized instrumentation, technical complexity, and ion-beam-induced damage^19,20^. In addition, vitrification of thicker specimens often requires high-pressure freezing or cryoprotectants, introducing further technical challenges and potential physiological perturbations.

Lysate-based workflows provide a more accessible alternative, in which cells are lysed in solution before the clarified extract is applied to EM grids without further purification^21–23^. Recent studies have recovered diverse high-resolution ribosomal structures from such cellular extracts, highlighting the potential of lysate-based cryo-EM to access endogenous macromolecular complexes^24,25^. However, bulk-lysis workflows homogenize cellular contents before vitrification, introducing dilution and time delays that can disrupt native spatial organization, alter molecular interactions, and perturb physiological state distributions.

A complementary approach is mechanical cell disruption on support surfaces, including cryo-EM grids. Previous unroofing approaches using sonication, filter-paper blotting, or pressurized fluid streams generate plasma membrane patches and cortical material suitable for cryo-ET of local cellular architecture^26–30^. By design, these methods primarily preserve membrane-proximal contents and retain only a limited fraction of the cell interior, leaving much of the intracellular material unrepresented in the final specimen. These limitations motivate the development of approaches that broaden access to diverse cellular targets while remaining compatible with high-resolution single-particle cryo-EM.

Here, we develop cryoGO (on-<u>G</u>rid <u>O</u>pening <u>cryo</u>-electron microscopy), a simple and high-throughput strategy that mechanically opens cells directly on EM grids immediately before vitrification. By specifically optimizing the exposure and dispersion of cellular contents into thin, electron-transparent layers, cryoGO produces samples compatible with high-resolution cryo-EM while minimizing dilution and the interval between cell disruption and freezing. We show that cryoGO enables near-atomic structural analysis of diverse endogenous macromolecular assemblies across a broad molecular-weight range. Using ribosomes as a benchmark system, we resolve a wide spectrum of translational states whose population distributions closely resemble those observed in *in situ* samples, supporting preservation of near-native physiological landscapes. The rapid workflow enables time-resolved interrogation of cellular physiology, allowing acute cellular responses to be captured on timescales of seconds.

## Results

### The cryoGO workflow

The cryoGO workflow integrates on-grid cell culture, on-grid opening, vitrification, cryo-EM data collection, and data processing into a streamlined pipeline (Fig. 1). Mammalian cells are first seeded and attached directly on poly-lysine (poly-K) coated cryo-EM grids. Once they reach optimal grid coverage, we perform on-grid opening with three consecutive rounds of automated filter paper blotting. For experimental consistency and reproducibility, we use a standard commercial vitrification device, the Thermo Scientific Vitrobot, which allows controlled blotting parameters and environmental conditions^31^. The defined forceful blotting from both sides of the grid mechanically disrupts the plasma membranes while wicking away excess liquid and depositing the exposed cellular contents into a thin film onto the EM grid. The grid is then immediately plunged into liquid ethane for vitrification. Importantly, the entire on-grid opening-to-freezing process is completed within a few seconds, minimizing preparation-induced perturbations while enabling the interrogation of rapid, time-dependent changes in the macromolecular landscape. The resulting vitrified grids are compatible with existing pipelines for either cryo-ET or tilt-free single-particle cryo-EM data collection and processing, enabling high-resolution structural determination of diverse macromolecular targets.

**Fig. 1.**
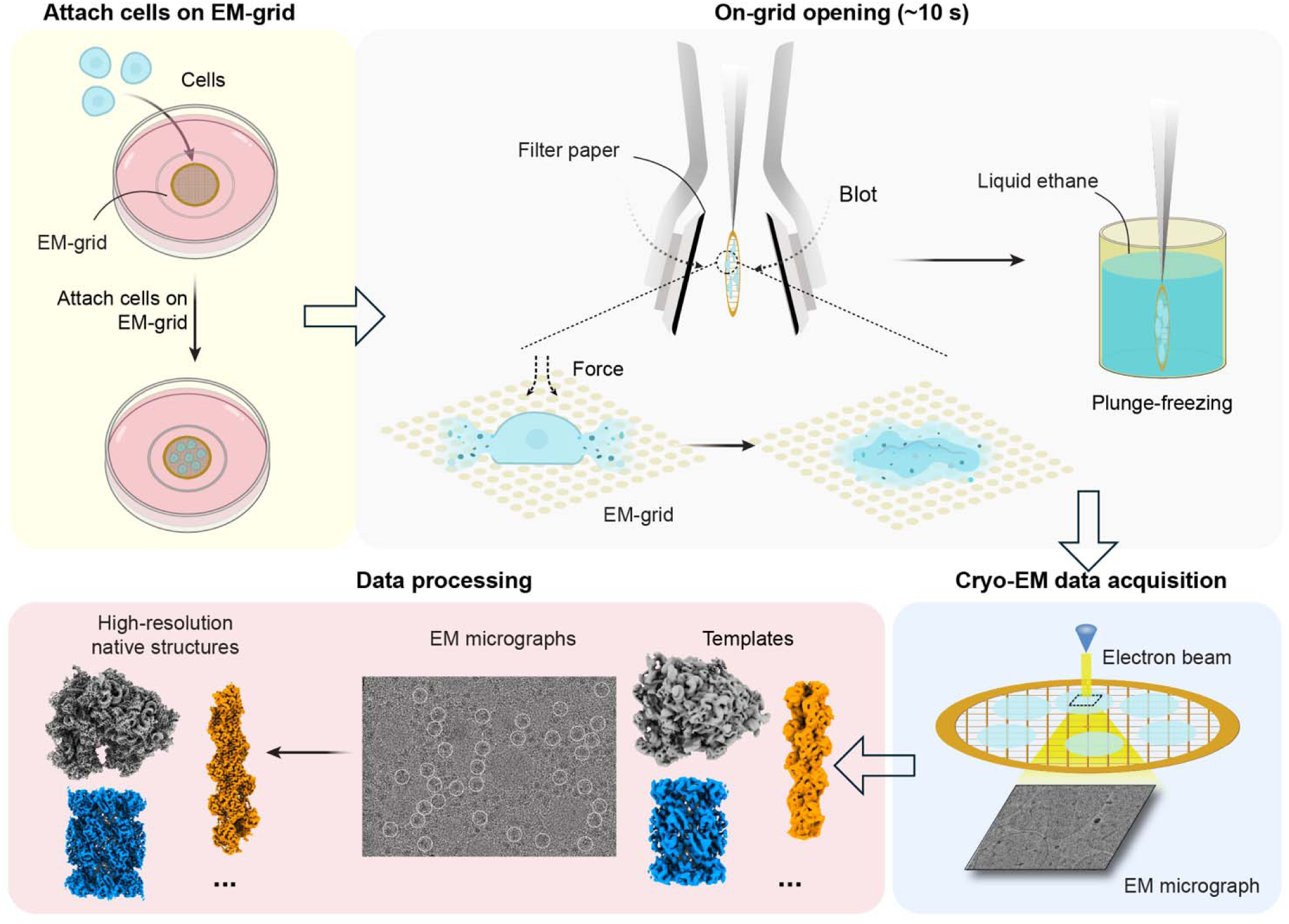
Schematic overview of cryoGO workflow. CryoGO pipeline integrates on-grid cell attachment, rapid on-grid opening and freezing, and structural determination. Cells are cultured directly on EM grids. Automated filter paper blotting applies mechanical force to open cells and spread the cellular contents into a thin film, followed by immediate plunge-freezing. High-contrast data of cellular contents are processed to resolve high-resolution native structures.

### On-grid opening spreads cellular contents into thin films

We first established and validated cryoGO using HEK293T cells, a widely used adherent human cell line. To validate the efficacy of on-grid opening, we used cryo-scanning electron microscopy (cryo-SEM) to compare the morphology of cryoGO samples to that of intact cells vitrified using the standard back-blotting method. In contrast to the bulky and convex morphology of intact cells, cryoGO samples formed a flattened, spread-out film (Fig. 2a). This physical disruption and spreading were also evident under cryo-transmission electron microscopy (cryo-TEM): intact cells are largely impenetrable to the electron beam, whereas cryoGO samples formed electron-translucent cellular spreads across grid squares (Fig. 2b). Consistently, quantitative gray-value measurements across intact-cell samples showed a sharp transition between electron-dense cells and empty grid regions. By contrast, cryoGO samples maintained intermediate gray values throughout the opened-cell region (Fig. 2c), indicating a continuous thin layer of cellular material. Furthermore, tomographic measurements of cross-sections confirmed that cryoGO samples formed a vitreous layer with a thickness of approximately 90 nm (Fig. 2d), well-suited for high-contrast single-particle cryo-EM imaging.

**Fig. 2.**
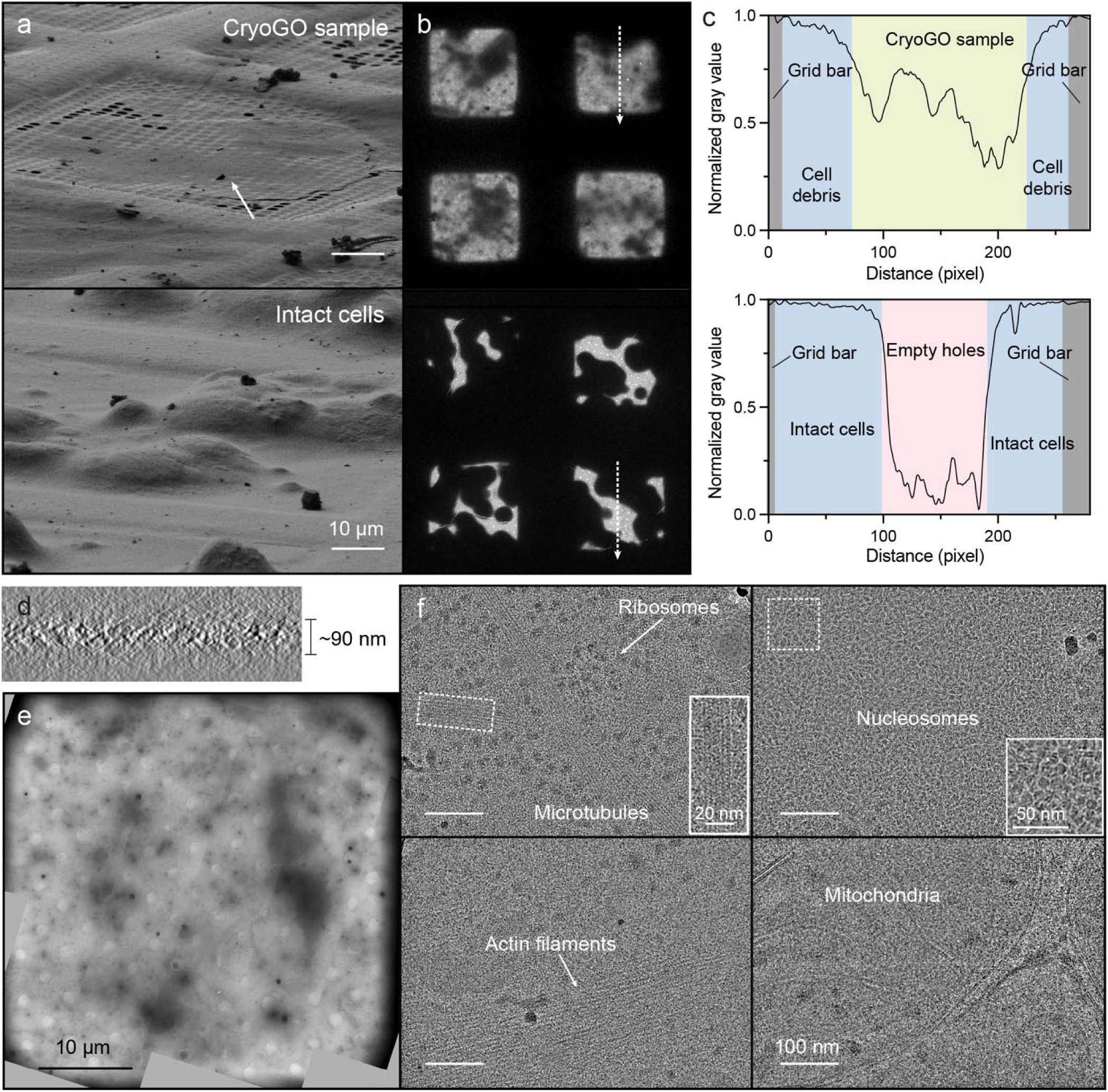
CryoGO disperses cellular contents into thin films for cryo-EM imaging. **a,** Cryo-SEM images showing the surface morphology of the spread film prepared by cryoGO (top) and intact cells prepared by back-blotting (bottom). Scale bars: 10 μm. **b**, Low-magnification cryo-EM atlas of cryoGO samples (top) and intact cells (bottom). **c**, Normalized image gray-value profiles along the dashed trajectories shown in **b**. **d**, Representative cross-sectional slice from a reconstructed tomogram of a cryoGO sample. **e**, Intermediate-magnification view of a representative grid square. Scale bar: 10 μm. **f**, Example cryo-EM micrographs revealing diverse cellular structures. A 5 Å resolution low-pass filter was applied. Scale bar, 100 nm. Insets show magnified views of the regions highlighted by dashed boxes containing microtubules (scale bar, 20 nm) and nucleosomes (scale bar, 50 nm).

### Systematic optimization of cryoGO sample preparation

To achieve consistent sample quality, we systematically investigated the effects of key preparation parameters. Evaluation of cell confluence revealed that sparse (∼20%) and moderate (∼50%) grid coverage supported efficient cell opening and formation of thin vitreous ice, whereas high confluence (∼80%) impeded disruption and resulted in clusters of intact cells unsuitable for imaging (Supplementary Fig. 1a, b). Consequently, all subsequent preparations were performed at ∼50% cell confluence, which provided the optimal balance of coverage, cell opening, and ice quality.

We next optimized the Vitrobot-controlled blotting conditions that drive the cell opening, specifically blot force and blot time. We observed that blot-force settings near 0 or a short blot time (1 s) were insufficient to disrupt the plasma membranes, leaving the majority of cells intact (Supplementary Fig. 1c, d). Conversely, excessive force (blot force 20) resulted in uneven sample distribution, potentially because angular compression of the blotting pads generated nonuniform blotting pressure and a thickness gradient across the grid. To achieve an optimal balance, we selected a blot force of 10 applied for 3 seconds for all subsequent preparations, as this condition maximized the proportion of grid squares containing a uniform layer of cellular material (Fig. 2e, Supplementary Fig. 1c, d). Notably, blotting conditions, particularly blot force, can vary among instruments, necessitating instrument-specific calibration and optimization. Our reported blot forces are all based on standard Vitrobot calibration, where a blot force of 0 results in the blotting pads barely touching each other.

### CryoGO enables high-resolution structural determination for diverse cellular macromolecules

Cryo-EM micrographs acquired under optimized cryoGO conditions exhibited high contrast and preserved diverse subcellular features, including membrane-bound organelles, cytoskeletal filaments, and macromolecular complexes such as ribosomes and nucleosomes (Fig. 2f). These properties provided an opportunity to resolve a broad range of endogenous cellular assemblies from a single cryo-EM dataset. However, the molecular crowding, compositional heterogeneity, and background noise characteristic of cellular environments pose substantial challenges for particle detection, classification, and alignment.

We first evaluated the conventional *ab-initio* single-particle reconstruction workflow using the highly abundant 80S ribosome as a benchmark, but found it insufficient for reliable particle detection in crowded cellular environments (Supplementary Fig. 2a). We next evaluated a template-based heterogeneous refinement strategy that iteratively classifies particles against target and “junk” references, progressively enriching correctly assigned particles while removing false positives^32,33^. This approach enabled high-resolution reconstructions of abundant targets including 80S ribosomes and nucleosomes (Supplementary Fig. 2b, c). Nevertheless, its performance remained limited by the accuracy of the initial particle assignments, restricting reliable recovery of lower-abundance and lower-contrast complexes, such as the 20S proteasome (Supplementary Fig. 2d).

We therefore adopted high-resolution 2D template matching^34–37^ using GisSPA for particle detection and alignment (Fig. 3a). By exhaustively evaluating correlations between experimental images and projections generated from high-resolution templates, this approach provides enhanced detection performance in crowded and heterogeneous cellular environments. To assess potential template bias, we employed an omit-map strategy^38^ in which selected regions were removed from the search template. The successful recovery of these omitted features, together with local refinements that converged to resolutions substantially exceeding those of the search templates, demonstrated that the final reconstructions were driven by experimental data rather than template propagation.

**Fig. 3.**
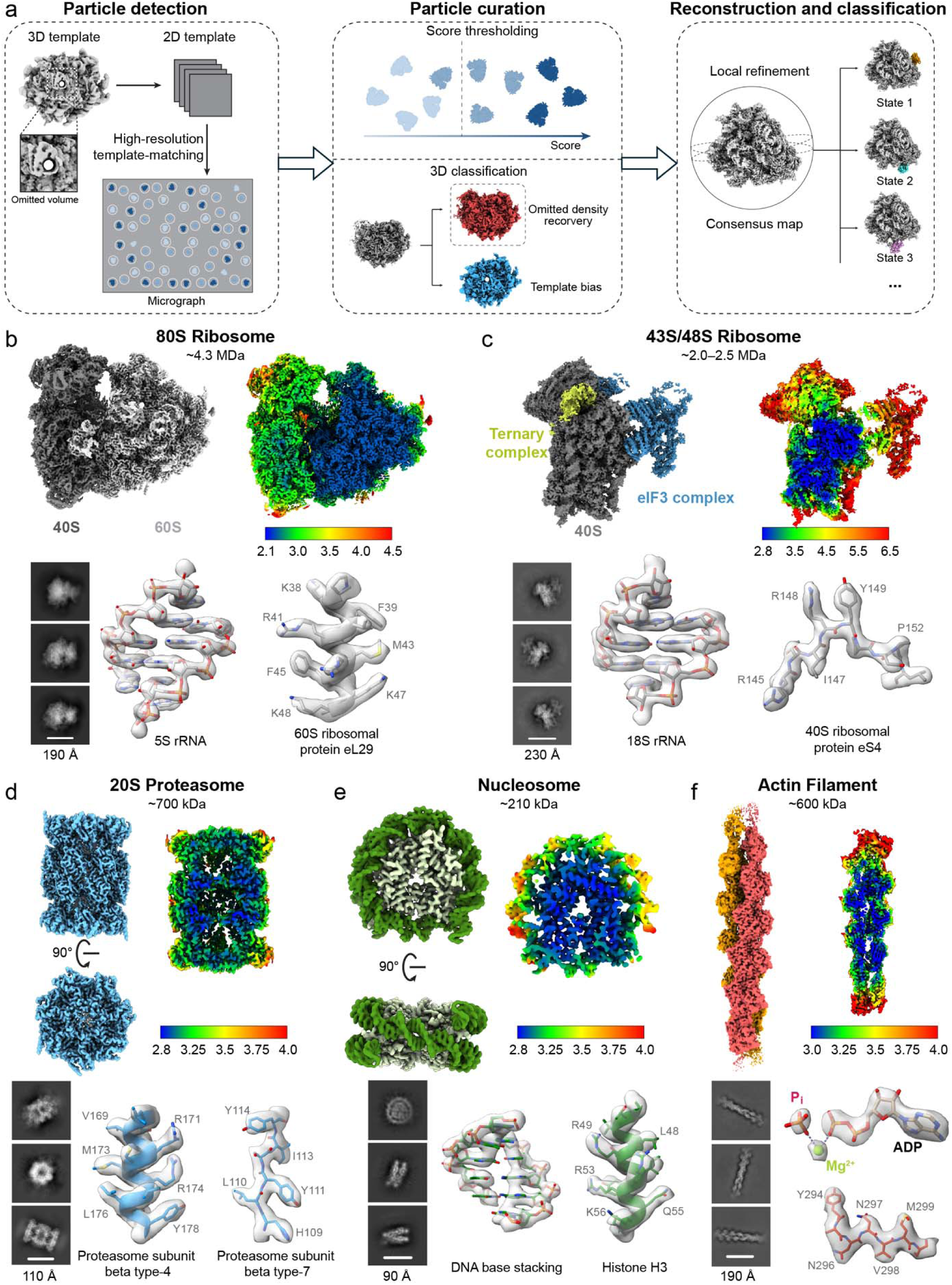
High-resolution structure determination of diverse cellular macromolecules using cryoGO. **a**, Overview of the data processing pipeline. Left: particle detection with high-resolution 2D template-matching algorithms using templates with omitted regions to minimize bias. Middle: particle curation through score thresholding and 3D classification to select high-quality particles. Right: reconstruction of consensus maps by local refinement and focused 3D classification to resolve various functional states. **b–f**, High-resolution structures of various macromolecules by cryoGO, including 80S ribosome (b), 43S/48S ribosome (c), 20S proteasome (d), nucleosome (e), and actin filament (f). For each example, top left: the consensus map; top right: local resolution map; bottom left: representative 2D class averages; and bottom right: representative high-resolution atomic model fitted into the cryo-EM map.

This strategy substantially improved particle detection (Supplementary Table 1) and enabled structural determination of diverse endogenous assemblies spanning a broad range of abundance, size, and functional states (Fig. 3b–f and Supplementary Fig. 3–4). The human 80S ribosome was resolved to a Nyquist-limited resolution of 2.2 Å, while nucleosomes were refined to 3.2 Å. We additionally resolved the transient 43S/48S translation initiation complex (2.8 Å), the 20S proteasome (3.0 Å), and extended actin filaments (2.9 Å). Across these diverse targets, the resulting maps exhibited well-resolved secondary-structure elements, amino-acid side chains, nucleotide bases, and ligand densities, demonstrating that cryoGO supports near-atomic structural analysis of a wide spectrum of endogenous cellular assemblies.

This ability to resolve such diverse targets relies on the improved particle contrast and signal-to-noise ratio afforded by cryoGO sample preparation of cellular specimens. To quantify this advantage, we reprocessed a previously published cryo-FIB-milled lamellae dataset^15^ using an identical computational workflow (Supplementary Fig. 5). Ribosome particles from cryoGO micrographs exhibited consistently higher template-matching cross-correlation scores than particles from cryo-FIB-milled samples, indicating improved particle signal quality (Supplementary Fig. 6a and Supplementary Table 2). Consistently, Rosenthal–Henderson analysis^39^ revealed a substantially lower apparent B-factor for the cryoGO dataset (84 Å^2^) than for the FIB-milled dataset (139 Å^2^) (Supplementary Fig. 6b), consistent with superior preservation of high-resolution information. Together, these analyses provide insight into how cryoGO supports high-resolution structure determination across a broad spectrum of endogenous cellular assemblies.

### CryoGO captures an extensive conformational ensemble of the translational machinery

Cellular macromolecules are inherently dynamic, a property essential for their functions. To assess whether cryoGO preserves the compositional and conformational landscape of endogenous cellular machines, we used the highly dynamic translational machinery as a test case. Through extensive 3D classification of ribosomal particles from cryoGO samples of human HEK293T cells, we resolved a diverse ensemble of 80S ribosome and 60S subunit states (Fig. 4 and Supplementary Fig. 7).

**Fig. 4.**
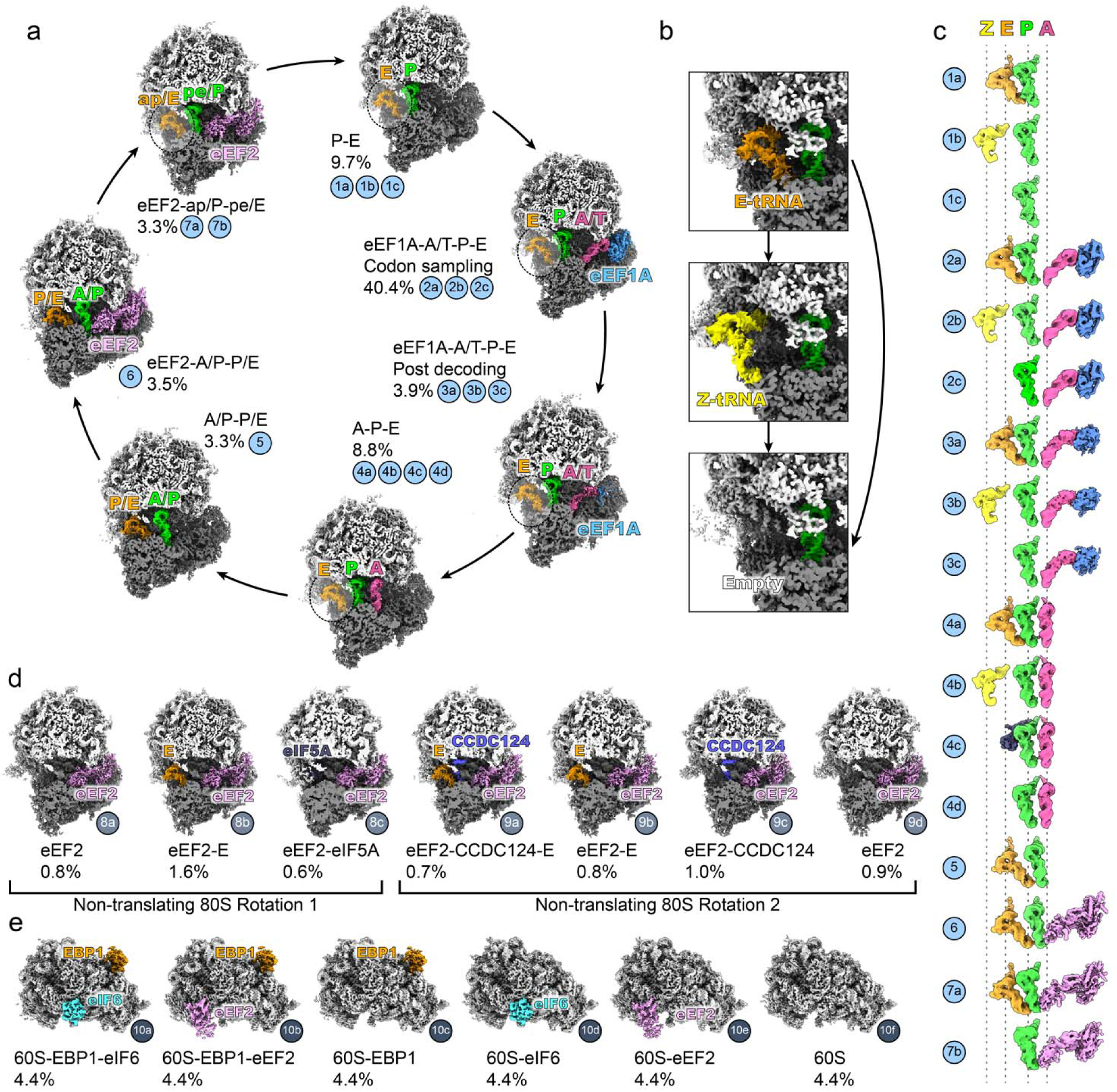
CryoGO captures an extensive conformational ensemble of the translational machinery. **a**, Cryo-EM maps of actively translating 80S ribosomes capturing sequential stages of the elongation cycle, clipped to show tRNAs and translation elongation factors. The percentages indicate the relative proportion of particles in each specific class. **b**, Magnified views of the E-site tRNA exit region (corresponding to the dashed circles in a), illustrating three distinct compositional states observed during tRNA dissociation: the canonical E-tRNA-bound state, the alternative Z-tRNA-bound state, and the empty state. **c**, Side-by-side structural comparison isolating the spatial trajectories of the tRNAs and translation elongation factors for each corresponding class in a. **d**, Cryo-EM maps of non-translating 80S ribosomes categorized into two distinct classes (Rotation 1 and Rotation 2) based on the relative rotation angle between the 40S and 60S subunits. **e**, Cryo-EM maps of the resolved 60S ribosomal subunits bound by various factors. For all figures, the 60S subunit is shown in light gray, the 40S subunit in dark gray, tRNA in A, P, E, and Z site in pink, green, orange, and yellow, respectively, eEF1A in sky blue, eEF2 in light magenta, CCDC124 in dark blue, eIF6 in cyan, and eIF5A in dark navy.

Analysis of the cryoGO data captured a trajectory across the entire eukaryotic elongation cycle (Fig. 4a–c). This trajectory included multiple states within each major elongation stage: pre-decoding ribosomes characterized by P-site tRNA occupancy and a vacant A-site (states 1a–1c), initial delivery of aminoacyl-tRNA to the A/T site and codon sampling/decoding by eEF1A (states 2a–2c, 3a–3c), accommodation of tRNAs into the A- and P-sites (states 4a–4d), and pre-translocation rotation with hybrid A/P-P/E tRNA configuration (state 5). We also captured eEF2 binding (state 6) and the progressive translocation toward the chimeric ap/P and pe/E positions (states 7a–7b). In parallel, focused comparison of E-site densities revealed a progressive E-site tRNA dissociation trajectory, including E-tRNA (states 1a, 2a, 3a, 4a and 7a), Z-tRNA (states 1b, 2b, 3b and 4b), and empty E-site configurations (states 1c, 2c, 3c, 4d and 7b) across multiple elongation intermediates (Fig. 4b, c).

In addition to translating ribosomes, we also resolved multiple non-translating 80S states distinguished by different degrees of 40S subunit rotation and association with specific regulatory factors, including eIF5A, eEF2, and CCDC124 (states 8a–8c, 9a–9d) (Fig. 4d). We further identified diverse populations of isolated 60S subunits bound by various factors, such as EBP1, eIF6, and eEF2 (states 10a–10f) (Fig. 4e). Together, these results demonstrate that cryoGO preserves a rich spectrum of endogenous ribosomal states and compositional heterogeneity.

### CryoGO preserves the near-native translational landscape

To further validate that the ribosomal state distributions captured by cryoGO faithfully reflect native translational landscapes, we compared the results with those obtained using two alternative sample-preparation methods (Fig. 5a): our previously published *in situ* data from FIB-milled lamellae^15^, which preserves the native cellular environment, and data from a bulk lysate sample prepared by the detergent extraction method^25^. To enable controlled comparison with the lamellae dataset, we matched the cell type and handling conditions across all samples, using the same HEK293A cell line and a 15-minute 5% glycerol treatment. To reduce computational uncertainty, particles from all three datasets were picked using identical 2D template-matching parameters and pooled into a combined stack for 3D classification (Supplementary Figs. 5 and 7). The entire classification pipeline was performed in three independent replicates with different random seeds to assess the reproducibility of state assignments under stochastic variation in the classification procedure. Quantitative analysis of the resulting populations revealed that the state distribution obtained *via* cryoGO closely resembled the native landscape observed in FIB-milled cells and maintained similar proportions of translating 80S, non-translating 80S, and 60S subunits (Fig. 5a, b, Supplementary Table 3). Minor discrepancies between the cryoGO and FIB-milling profiles may have arisen from inherent biological variation or perturbations during sample preparation. By contrast, the bulk lysate sample exhibited severe deviations marked by an artifactual accumulation of eIF5A-bound non-translating ribosomes (state 8c), accompanied by a substantial depletion of several translating ribosomal states (states 1b, 2b, 3b, 4b, and 6) (Fig. 5a).

**Fig. 5.**
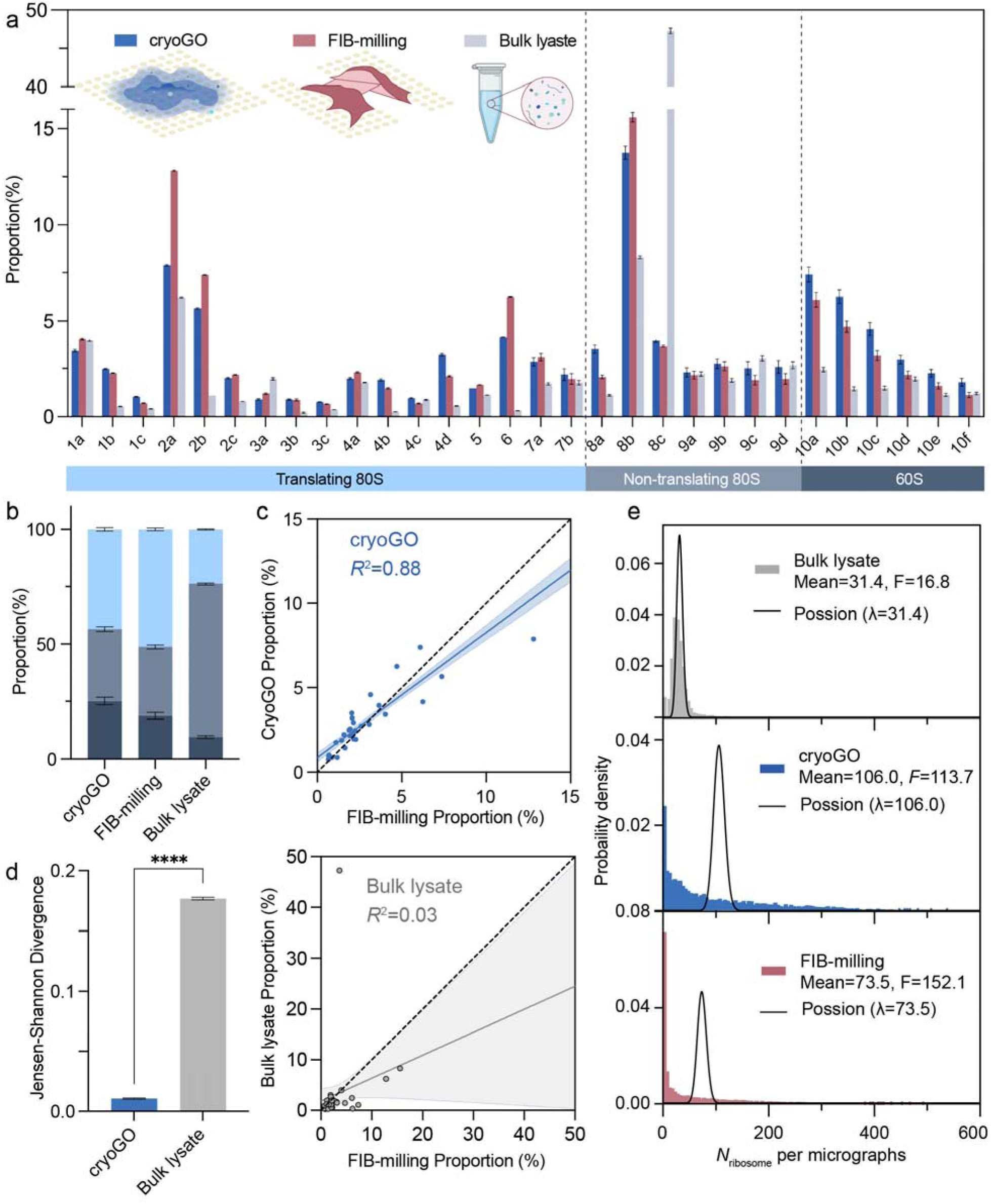
CryoGO preserves the near-native translation landscape and partial cell-derived spatial heterogeneity. **a**, Detailed comparison of ribosome state distributions obtained *via* cryoGO (blue), FIB-milling (red, which serves as a native-state reference), and bulk lysate (grey). **b**, Stacked bar chart summarizing the proportions of translating 80S, non-translating 80S, and 60S subunits across the three methods. **c**, Linear regression analyses comparing the state proportions of cryoGO (top) and bulk lysate (bottom) against the FIB-milling baseline. The dashed line represents the identity line (y = x), and the solid lines represent the best linear fits. The shaded regions indicate the 95% confidence intervals of the regression lines. **d**, Jensen-Shannon divergence scores quantifying the dissimilarity of the ribosome state distributions obtained *via* cryoGO and bulk lysate relative to the FIB-milling native-state reference. \*\*\*\**p* < 0.0001. For **a**, **b**, and **d**, data are presented as mean ± standard deviation derived from the three independent classification analyses. **e,** Per-micrograph ribosome count distributions for the bulk lysate, cryoGO, and FIB-milled samples. The solid black curve is the theoretical Poisson distribution computed at the sample mean.

This high fidelity was further corroborated by statistical analyses. Linear regression demonstrated a strong correlation between the state distributions of cryoGO and the *in situ* FIB-milling reference (*R*² = 0.88), whereas the bulk lysate result showed virtually no correlation (*R*² = 0.03) (Fig. 5c, Supplementary Table 4). Furthermore, we evaluated the Jensen-Shannon (JS) divergence, a standard symmetric measure of distributional dissimilarity^40^. CryoGO exhibited an exceptionally low JS divergence from the FIB-milling standard, in contrast to the markedly elevated divergence for the bulk lysate (Fig. 5d, Supplementary Table 4). Collectively, these metrics support the conclusion that cryoGO preserves the translational landscape at a level comparable to that of FIB-milled *in situ* samples and can serve as an effective approach for probing near-native molecular ensembles.

### CryoGO partially retains cell-derived spatial heterogeneity

The spatial context of a macromolecule fundamentally shapes its conformational and compositional state. To evaluate the extent to which the cryoGO workflow preserves the spatial heterogeneity of the cell, we analyzed the per-micrograph ribosome count (*N*_ribosome_) distributions across samples generated by the three different preparation methods (Fig. 5e). Theoretically, for homogeneously mixed independent particles (count density λ), *N*_ribosome_ follows a Poisson distribution with a variance-to-mean ratio (Fano factor, *F*) of 1 (Fig. 5e). In the bulk lysate sample, in which the cellular contents were homogenized in solution, the distribution was approximately unimodal and Poisson-like but exhibited overdispersion (*F*=16.8), potentially owing to residual polysome clustering and other sources of heterogeneity. By contrast, cellular samples prepared by FIB-milling showed extreme overdispersion (*F*=152.1) and a strongly right-skewed, long-tailed distribution, consistent with the heterogeneous spatial organization of ribosomes in cells. Notably, the cryoGO sample also displayed severe overdispersion (*F*=113.7) with a long-tailed distribution reminiscent of that of the FIB-milled sample (Fig. 5e). Across all evaluated distribution metrics, the cryoGO profile was intermediate between those of the bulk lysate and FIB-milled samples (Supplementary Table 5). Similar distribution characteristics were also observed across all other evaluated cryoGO datasets (Supplementary Fig. 8). This clear departure from random mixing indicates that cryoGO does not fully homogenize cellular contents and partially preserves cell-derived spatial heterogeneity.

### CryoGO captures time-resolved dynamic responses to physiological perturbations

By coupling cell opening and vitrification into a single, rapid procedure, cryoGO offers temporal resolution on the scale of seconds, facilitating the capture of progressive or transient cellular responses. To demonstrate that cryoGO can track biological perturbations over a time series, we first investigated the translational stress response to nutrient depletion. We subjected HEK293T cells to amino acid starvation (–aa) for varying durations (1 hr, 8 hr, and 24 hr) (Fig. 6a), and the resulting cryoGO datasets revealed time-dependent remodeling of the cellular translation landscape. As expected, compared to the unperturbed baseline, amino acid starvation led to a marked reduction in the population of actively translating ribosomes and a concomitant increase in non-translating 80S complexes (Fig. 6b, c, Supplementary Fig. 9a). Specifically, we observed a progressive decline in the proportion of translating ribosomes from 1 hr to 8 hr, reflecting the suppression of global protein synthesis triggered by severe nutrient depletion^41^. After 24 hr of starvation, the proportion of translating ribosomes exhibited a modest rebound, which may reflect nutrient recycling during prolonged starvation^42,43^ (Fig. 6c).

**Fig. 6.**
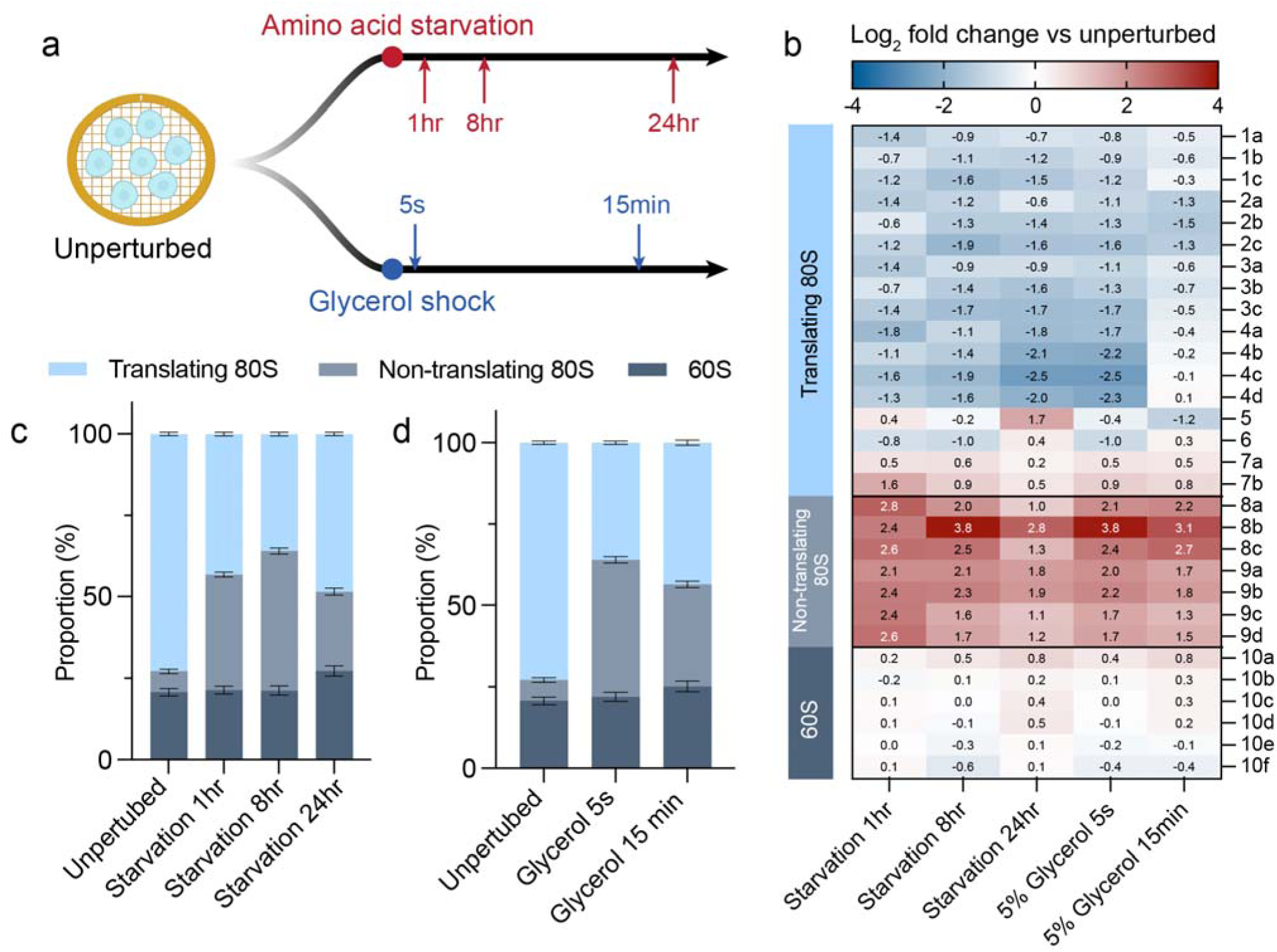
CryoGO captures time-resolved translational dynamics in response to physiological perturbations. **a**, Schematic representation of the experimental design. Unperturbed cells cultured on grids were subjected to either progressive amino acid starvation (sampled at 1 hr, 8 hr, and 24 hr) or 5% glycerol shock (sampled at 5 s and 15 min). **b**, Heatmap illustrating the observed relative population shifts of ribosomal states across the perturbation time courses. Values represent the log_2_ fold change for each state proportion compared to the unperturbed baseline. Blue indicates depletion and red indicates enrichment. **c, d,** Stacked bar charts summarizing the redistributions of translating 80S, non-translating 80S, and 60S pools during amino acid starvation (**c**) and glycerol shock (**d**). Data are presented as mean ± standard deviation derived from the three independent classification analyses.

Having established the ability to track progressive cellular responses over extended time courses, we next evaluated whether cryoGO could capture acute cellular responses on timescales of seconds. To test this, we exposed HEK293T cells to a rapid shock of 5% glycerol for approximately 5 seconds (Fig. 6a). Even this brief exposure was sufficient to induce a substantial decrease in actively translating ribosomes, shifting the translational landscape toward the severely perturbed state observed after 15 minutes of 5% glycerol treatment (Fig. 6b, d, Supplementary Fig. 9b). The observed perturbation after sub-minute exposure to standard cryoprotectants highlights a potential caveat of cryoprotectant-dependent workflows, a limitation that cryoGO avoids by coupling on-grid opening directly to vitrification without cryoprotectant treatment. Collectively, these findings support the use of cryoGO for time-resolved analysis of cellular responses across a broad range of timescales, from prolonged adaptation to rapid stress responses.

### CryoGO is broadly compatible with diverse cellular systems and requires minimal sample material

To establish the generalizability of the cryoGO workflow, we extended the method to a diverse panel of additional cellular specimens, including adherent cell lines (human colon carcinoma RKO and human gastric adenocarcinoma AGS cells), suspension cell lines (insect ovary Hi5 and human T-cell leukemia MT-4 cells), virus-infected cells (HIV pseudovirus infected human embryonic kidney HEK293T cells^44^ and MT-4 cells), and specialized primary cells (mouse oocytes) (Supplementary Fig. 10). Despite their distinct sizes and growth properties in standard culture, all cells readily attached to poly-K-coated EM grids and were efficiently prepared with the cryoGO workflow. Cryo-EM micrographs demonstrated the capture of cell-specific structural features, including active HIV virion production and budding in infected cells and cytoplasmic lattice assemblies unique to mammalian oocytes^45^ (Supplementary Fig. 10). Notably, cryoGO requires only small amounts of starting material for high-resolution structure determination, ranging from a few microliters of cell suspension to cells grown directly on an EM grid or ∼6 oocytes deposited on a grid^45^.

Ribosome-state analysis of the HIV-infected MT-4 dataset revealed a prominent population of non-translating ribosomes bound by eIF5A (state 8c) (Supplementary Fig. 9c), whereas the corresponding fraction was very small in HEK293T cells, consistent with the elevated expression of eIF5A reported in immune cell lineages^46^. The ability to systematically detect such lineage-specific structural and compositional features underscores the broad applicability of cryoGO across diverse cell types, growth formats, biological states, and cells of different sizes.

## Discussion

Inspired by the concept of on-grid unroofing^26–30^, cryoGO broadens access beyond membrane-proximal material by optimizing cell opening for whole-cell content recovery. It enables near-atomic-resolution reconstruction of diverse endogenous assemblies through a mechanistically simple operation: blot-force-driven spreading converts thick cells into a nanometer-scale film that allows electron transmission while preserving near-native structural information. The resulting thin, particle-rich specimens provide high image contrast and improved signal-to-noise relative to cryo-FIB-milled lamellae, supporting more efficient particle detection and higher-resolution reconstructions. The approach extends to targets that have been difficult to visualize in native cellular environments, including low-abundance complexes, modestly sized assemblies, and transient functional intermediates. The ability to resolve these targets without purification or the technical demands of cryo-FIB-milling underscores the practical accessibility that distinguishes cryoGO from existing approaches.

Using ribosomes as a benchmark, we observed close agreement of translational state distributions between cryoGO and FIB-milling samples, our current best proxy for the native cellular state, indicating that rapid on-grid opening preserves much of the native conformational and compositional landscape. Two features of the workflow likely contribute to this fidelity: the seconds-scale interval between cell opening and vitrification helps preserve transient translational intermediates, and on-grid mechanical disruption avoids the dilution and complete bulk homogenization introduced by lysate extraction. In our hands, lysate extraction substantially altered state distributions. Quantitative analyses further revealed partial retention of cell-derived spatial heterogeneity. Although much of the native cellular architecture is inevitably lost as a three-dimensional cell is flattened into a nanometer-scale layer, our results suggest that cryoGO retains sufficient local organization to reveal biologically meaningful cellular state distributions.

By integrating cell opening with immediate vitrification, cryoGO also provides temporal resolution on the scale of seconds. This capability enabled capture of both long-term cellular adaptation to nutrient deprivation and rapid responses to acute cryoprotectant stress. Notably, we found that even brief exposure to a commonly used cryoprotectant was sufficient to induce substantial translational perturbations, underscoring the importance of minimizing sample-preparation-induced artifacts in native cryo-EM studies. More broadly, cryoGO is readily compatible with a wide range of upstream cellular treatments, enabling time-resolved analysis of endogenous molecular machines under diverse physiological conditions.

The successful application of cryoGO across diverse cell types, growth formats, and biological states underscores its versatility and accessibility. Together with its ability to generate complete datasets from a small number of cells, this broad compatibility opens opportunities to determine the high-resolution structures of endogenous macromolecules from specialized, limited-availability, and physiologically complex samples. Because the method requires only standard cryo-EM grids and a routine commercial vitrification device, it lowers the technical barrier for high-resolution native cellular structural biology and provides a practical route for connecting endogenous molecular structures with cell-specific physiology.

## Methods

### Cell culture and maintenance

Human HEK293T, HEK293A, RKO, and AGS cells were cultured in Dulbecco’s modified Eagle medium (DMEM) supplemented with 10% (v/v) fetal bovine serum (FBS) and 1% penicillin–streptomycin. Cells were maintained in a humidified incubator at 37 °C with 5% CO and passaged every 2–3 days before reaching 70–80% confluence.

MT-4 cells were cultured in RPMI-1640 medium supplemented with 10% (v/v) FBS and 10 μg/ml gentamicin. Cells were maintained at 37 °C in a humidified incubator with 5% CO_2_ and passaged every 2–3 days to maintain a density between 2×10^5^ and 1×10^6^ cells/mL. Insect Hi5 cells were maintained in ESF 921 medium supplemented with 10 μg/ml gentamicin. Cells were cultured at room temperature in an ambient-air incubator with continuous shaking and passaged regularly to maintain healthy suspension growth. Mouse oocytes were isolated and matured as described previously^45^. Prophase I-arrested oocytes were isolated from the 6–8-week-old CD1 female mice and matured *in vitro* into MII. All animal procedures were approved by the Yale University Institutional Animal Care and Use Committee (protocols 2024-20408 and 2024-20562).

### CryoGO sample preparation

R2/1 holey carbon gold grids (200 mesh, Quantifoil) coated with a 2-nm continuous carbon film, or R2/4 holey carbon gold grids (200 mesh, Quantifoil), were glow-discharged at 20 mA for 25 s. For sterilization, the grids were submerged in 70% ethanol and exposed to ultraviolet (UV) irradiation for 1 hour. Following sterilization, the grids were washed and incubated in a 0.01% poly-lysine solution for 1 hour to promote cell attachment. Prior to cell seeding, the coated grids were thoroughly washed five times with phosphate-buffered saline (PBS) to remove residual poly-lysine. Cell suspensions were subsequently applied to the grids, targeting an optimal confluence of approximately 50%, unless otherwise indicated. For standard cell lines, cells were generally seeded in 35-mm dishes with 2 mL of a 5 × 10^5^ cells/mL suspension, with exact seeding conditions empirically optimized depending on the specific cell type. The seeded grids were then incubated under standard culture conditions until stable cell attachment was achieved. For primary cell preparations, 5 to 7 mouse oocytes were manually transferred onto each grid. The cell-bearing grids were then mounted in a Thermo Scientific Vitrobot operated at 10 °C and 100% humidity. On-grid opening was performed through three consecutive rounds of automated filter-paper blotting. Unless otherwise stated, the optimized condition consisted of a blot force of 10 with successive blot times of 1 s, 1 s, and 3 s. Immediately following the final blot, the grids were plunge-frozen in liquid ethane for vitrification.

### Optimization of on-grid opening

HEK293T cells were used to systematically optimize the key parameters of the on-grid opening. To determine the optimal cell density, cells were seeded and cultured on EM grids to achieve target confluences of approximately 20%, 50%, and 80%. The confluence of the attached cells was evaluated using light microscopy. Blot-force optimization was performed with blot force 0, 10 and 20 while keeping the blot time fixed at 3 s. Blot-time optimization was performed with 1, 3 and 5 s blot times while keeping the blot force fixed at 10. The resulting grids were evaluated *via* cryo-EM to assess the efficiency of cell opening.

### Cellular stress conditions and viral infection

For amino acid starvation (–aa), HEK293T cells cultured on EM grids were washed three times with PBS and incubated in an amino acid-depleted medium for 1, 8, or 24 h prior to cryoGO preparation. For acute glycerol stress, EM grids bearing HEK293T cells were submerged in culture medium supplemented with 5% (v/v) glycerol for ∼5 s and immediately processed *via* the cryoGO workflow. For matched comparisons with the published cryo-FIB-milled lamellae data^15^, grid-cultured HEK293A cells were incubated in a medium containing 5% (v/v) glycerol for 15 min under standard culture conditions prior to cryoGO preparation. For viral infection experiments, MT-4 cells were infected with a VSV-G-pseudotyped, envelope-deficient HIV-1 pseudovirus at a multiplicity of infection of 2.5. At 6 h post-infection, the infected cells were seeded and cultured onto EM grids and subsequently processed *via* the cryoGO workflow. HIV-1G- or HIV-3G-infected HEK293T cells were kindly provided by the Wei-Shau Hu laboratory, with viral infections performed according to previously established protocols^44^. These infected cells were subsequently prepared using the cryoGO workflow as described.

### Bulk-lysate sample preparation

Bulk-lysate samples were prepared following previously published protocols^25^. Briefly, HEK293A cells were seeded and cultured in 150-cm^2^ plates until reaching confluence. For glycerol treatment, cells in two plates were incubated for 15 min after the culture medium was replaced with medium supplemented with 5% (v/v) glycerol. Following the treatment, the cells were washed twice with PBS, detached using Trypsin-EDTA, and harvested by centrifugation at 300 × *g* for 5 minutes. The resulting cell pellets were resuspended in 300 μL of ice-cold semi-permeabilization buffer (25 mM HEPES pH 7.2, 110 mM KOAc, 15 mM Mg(OAc)_2_, 1 mM DTT, 0.015% digitonin, 2× protease inhibitor cocktail, 40 U/mL RNase inhibitor, and 1 mM EGTA) and incubated at 4 °C for 5 minutes. The lysate was then clarified by centrifugation at 1,000 × *g* for 5 minutes at 4 °C, and the resulting supernatant was collected for downstream grid preparation. For vitrification, 5 μL of the clarified lysate was applied onto glow-discharged R2/1 holey carbon copper grids (300 mesh, Quantifoil). The grids were mounted in a Thermo Scientific Vitrobot at 10 °C and 100% relative humidity, blotted for 6.5 s with a blot force of 1, and immediately plunge-frozen in liquid ethane.

### Cryo-SEM sample preparation and imaging

CryoGO samples of HEK293T cells were prepared as described above. For intact cell controls, HEK293T cells were similarly grown and attached onto EM grids. Grids were mounted in a Vitrobot at 10 °C, 100% relative humidity, manually blotted from the back side for 4 s, and immediately plunge-frozen in liquid ethane. Sample morphology was subsequently evaluated using an Aquilos cryo-FIB/SEM system (Thermo Fisher Scientific). Prior to imaging, the grids were sputter-coated with an organometallic platinum layer for 30 s under cryogenic conditions.

### Cryo-electron tomography data acquisition and processing

Cryo-ET data of HEK293T cryoGO samples were acquired using a 200 kV Glacios electron microscope (Thermo Fisher Scientific) equipped with a Gatan K3 direct electron detector. Tilt series were collected at a magnification of 45,000×, corresponding to a physical pixel size of 0.86 Å/pixel. A dose-symmetric tilt scheme was employed, covering an angular range from −54° to +54° with a 3° increment. Tomography data processing and 3D reconstruction were performed using the EMAN2 software suite^47^. The reconstructed tomograms were used to measure the cross-sectional thickness of the cryoGO sample.

### Cryo-EM data collection and image preprocessing

Single-particle cryo-EM datasets were acquired from electron-translucent regions in cryoGO samples or bulk-lysate samples. Unless otherwise indicated, data were collected on a 300 kV Titan Krios electron microscope (Thermo Fisher Scientific) equipped with a Gatan BioQuantum energy filter (20-eV slit width) and a K3 direct electron detector (Gatan). Images were recorded at a nominal magnification of 81,000× (physical pixel size of 1.068 Å/pixel) with a total electron dose of 50 e /Å² and a target defocus range of −0.5 to −1.5 μm. For –aa samples, datasets were collected on a 200 kV Glacios electron microscope (Thermo Fisher Scientific) equipped with a K3 detector at 45,000× magnification (physical pixel size of 0.86 Å/pixel) using a total electron dose of 50 e /Å^2^. Automated data collection was performed using SerialEM^48^. The acquired movies were processed for motion correction^49^ and contrast transfer function (CTF) correction^50^ using cryoSPARC^51^. Micrographs with poor ice quality, strong drift, and severe contamination were excluded.

### Conventional *ab-initio* and heterogeneous refinement data processing workflows

A conventional *ab-initio* processing workflow for the 80S ribosome was applied to the unperturbed HEK293T cell dataset. Particles were initially detected using the template picker in cryoSPARC with a human 80S ribosome reference (EMD-71440), extracted with a 512-pixel box (binned to 256 pixels), and subjected to 2D classification, *ab-initio* reconstruction, and heterogeneous refinement.

For the referenced heterogeneous refinement strategy, the cryoSPARC-picked 80S ribosome particles were split into four subsets and subjected to three rounds of heterogeneous refinement, without prior 2D classification, against the target reference and three “junk” references created by low-pass filtering the refence at 100 Å. The particles refined to high resolution were retained and re-extracted unbinned, followed by global and local CTF refinements and local refinement. For the nucleosome, particles were selected using the cryoSPARC template picker with a nucleosome reference (EMD-32220), extracted with a 200-pixel box (binned to 80 pixels), and subjected to 13 rounds of heterogeneous refinement with C2 symmetry against the target and three junk references, followed by unbinned re-extraction, CTF refinements, and local refinement. For the 20S proteasome, particles were similarly selected using the cryoSPARC template picker (EMD-4877), extracted with a 256-pixel box, and subjected to seven rounds of heterogeneous refinement with C2 symmetry against the target and four junk references. Full processing details for these workflows are summarized in Supplementary Fig. 2.

### 2D template matching data processing workflows

Initial particle detection was performed using the GisSPA algorithm^37^. Target templates were low-pass filtered and specific structural volumes were omitted prior to the search. Unless otherwise stated, template matching was executed on bin-3 micrographs with a 3° angular step for both in-plane and out-of-plane rotations. Initial particles were filtered using normalized cross-correlation score thresholds (ranging from 6.2 to 6.6) optimized per dataset. Particles were then imported into RELION^52^ and cryoSPARC alongside their GisSPA-assigned orientations for extraction and subsequent analysis.

For 60S/80S ribosomes, a human 60S reference (EMD-71413, low-pass filtered to 6.4 Å, then resampled to 3.2 Å/pixel with a cylindrical volume removed) was used as the search template. Particles were extracted with a 432-pixel box using RELION^52^ and subjected to local refinement, global and local CTF refinements, and 3D classification utilizing a 60S focus mask in cryoSPARC. Additional false positives were removed *via* per-particle scale filtering. This processing pipeline was systematically applied to all datasets, including the bulk lysate, the previously published FIB-milled lamellae data^15^, and the cryoGO datasets. Retained particles from each dataset underwent a further round of local refinement before being merged into a single combined stack for all subsequent classification and analysis. Processing details for all the datasets can be found in Supplementary Figs. 3 and 5.

For 43/48S ribosomes, an in-house human 48S template (low-pass filtered to 6.4 Å, then resampled to 3.2 Å/pixel with a selected RNA duplex volume omitted) was used to search the HEK293T cell dataset. Following extraction (bin-2), particles underwent local refinement, 3D classification, and per-particle scale filtering. Retained particles were re-extracted unbinned, followed by global and local CTF refinements, local refinement, and two additional rounds of 3D classification to separate the 43/48S and 40S classes (Supplementary Fig. 3). For the nucleosome, a human nucleosome template (EMD-47924, resampled to 2.1 Å/pixel with a cylindrical volume omitted) was used to search the HIV-infected MT-4 dataset on bin-2 micrographs with a 3° angular step. Particles underwent local refinement with C2 symmetry and 3D classification, followed by CTF refinement, an additional 3D classification, and a final local refinement. For the actin filament, an actin template (EMD-15108 resampled to 4.3 Å/pixel resolution with a selected volume omitted) was applied to the HIV-1 1G-infected HEK293T dataset on bin-4 micrographs with a 5° angular step. Extracted particles (bin 2) were processed *via* local refinement, 3D classification, and per-particle scale filtering. Retained particles were then re-extracted unbinned, subjected to CTF refinement, and locally refined. For the 20S proteasome (EMDB-4877, resampled to 2.1 Å/pixel), particles were searched on bin-2 micrographs with a 3° angular step across four datasets (HEK293T, HIV-1G- and HIV-3G-infected HEK293T cells, and HIV-infected MT-4 cells). Extracted particles (256-pixel box) were merged and subjected to local refinement, 3D classification, per-particle scale filtering, CTF refinements, and a final local refinement. Processing details for the nucleosome, actin filament, and 20S proteasome are in Supplementary Fig. 4. All the refinement statistics are in Supplementary Table 6. Fourier shell correlation (FSC) curves for all final reconstructions are in Supplementary Fig. 11.

### Particle-quality and Rosenthal-Henderson analyses

To compare data quality between datasets from cryoGO and FIB-milled samples, the distributions of the normalized cross-correlation scores derived from GisSPA searches for the final 60S/80S ribosome particles were compared. Rosenthal-Henderson plots were generated using the ResLog analysis tool in cryoSPARC by reconstructing randomly sampled particle subsets of increasing size. The squared spatial frequency at the reported resolutions was plotted against the natural logarithm of the particle numbers, and apparent B-factors were estimated from the slopes of the resulting linear fits.

### 3D classification for ribosome state-population analysis

60S and 80S ribosome particles from all independent datasets were pooled for combined analysis to reduce computational variations. An initial 3D classification, focused on 40S, separated 80S ribosomes by their major rotational states and isolated 60S subunits lacking 40S density. For the 60S subunits, two independent focused classifications were performed in parallel: one targeting the EBP1-binding region to determine EBP1 occupancy, and another targeting the A-site to identify eEF2 or eIF6 binding. Intersecting particles in these classes yielded six distinct 60S classes from all factor-binding combinations. Similarly, the 80S rotational classes were subjected to hierarchical and combinatorial classifications using focus masks on the A-, P-, and E-sites to resolve specific tRNA configurations and elongation factor-bound states. Detailed classification procedures are summarized in Supplementary Fig. 7.

Following classification, the final particle sets for each resolved state from the combined dataset were intersected with the original individual datasets to recover dataset-specific population statistics. To ensure statistical robustness, the entire 3D classification pipeline was performed in three independent replicates with randomly assigned seeds. Linear regression and Jensen-Shannon divergence analyses were used to compare state distributions between cryoGO, FIB-milling, and bulk lysate preparation methods. The refinement statistics are summarized in Supplementary Table 7. FSC curves for all resolved states from the HEK293T dataset are summarized in Supplementary Fig. 11.

### Per-micrograph ribosome particle count distribution analysis

For each sample preparation method (bulk lysate, cryoGO, and FIB-milling), the number of 60S/80S ribosome particles retained in the final reconstruction was recorded for each micrograph, defined as the per-micrograph ribosome particle count (*N*_ribosome_). From these per-micrograph counts, the empirical mean ⟨*N*⟩, variance σ2, Fano factor *F*=σ2/⟨*N*⟩, and relative variance CV^2^=(*F*-1)/⟨*N*⟩ were computed across all micrographs for each sample.

## Data availability

The cryo-EM density maps generated in this study have been deposited in the Electron Microscopy Data Bank (EMDB). The consensus density maps of human 80S ribosomes obtained by cryoGO from HEK293T cells, HEK293T cells treated with 5% glycerol for 5 s, HEK293A cells treated with 5% glycerol for 15 min, HEK293T cells subjected to amino acid starvation for 1 h, 8 h, or 24 h, and HIV-infected MT-4 cells have been deposited under accession codes EMD-77427, EMD-77458, EMD-77459, EMD-77462, EMD-77463, EMD-77464, and EMD-77465, respectively. The reprocessed human 80S ribosome map from the published HEK293A FIB-milled lamella dataset^15^ has been deposited under accession code EMD-77460. The human 80S ribosome map from HEK293A cells treated with 5% glycerol for 15 min and prepared by bulk lysate extraction has been deposited under accession code EMD-77461. The density maps of individual human 60S and 80S ribosomal states obtained by cryoGO from HEK293T cells have been deposited under accession codes EMD-77428 to EMD-77457. These correspond to states 1a, 1b, 1c, 2a, 2b, 2c, 3a, 3b, 3c, 4a, 4b, 4c, 4d, 5, 6, 7a, 7b, 8a, 8b, 8c, 9a, 9b, 9c, 9d, 10a, 10b, 10c, 10d, 10e, and 10f, respectively. The consensus density maps of the human 43S/48S ribosomal complex, nucleosome, 20S proteasome, and actin filament obtained by cryoGO have been deposited under accession codes EMD-77466, EMD-77467, EMD-77468, and EMD-77469, respectively. A complete list of accession codes is provided in Supplementary Tables 6 and 7. Source data are provided with this paper.

## Acknowledgements

We thank the David Haselbach lab at the Research Institute of Molecular Pathology for providing the RKO cell line, the Stavroula Hatzios lab at Yale University for providing the AGS cell line, and the Kai Zhang lab at Yale University for providing the HEK293A cell line. We thank Drs. Olga Nikolaitchik and Alice Duchon in the Wei-Shau Hu lab at the National Cancer Institute for preparing and providing HIV 1G/3G virus-infected HEK293T cells, and the Steven Tang lab at Yale University for providing the Hi5 cell line. We also thank Dr. Athanasios Bakasis and the Bieniasz–Hatziioannou lab at Rockefeller University for providing the MT-4 cell line and HIV pseudovirus, and the Binyam Mogessie lab at Yale University for providing mouse oocytes. We thank Yale CryoEM Resource facilities for support with cryo-EM data collection. We thank the Yale Center for Research Computing for support with computing resources. This work was supported by the National Institutes of Health through grant R01AI192025 and Yale discretionary funds to Y.X.

## Author contributions

Conceptualization: YX. Methodology: YL, YZ, CW, CZ, YX. Investigation: YL, YZ, CW, CZ. Visualization: YL, YZ, YX. Project administration: YX. Supervision: YX. Funding acquisition: YX. Writing – original draft: YL, YZ, YX. Writing – review & editing: YL, YZ, YX.

## Competing interests

The authors declare no competing interests.

## Notes

### Competing Interest Statement

The authors have declared no competing interest.

