## Supplementary Information for "CryoGO enables high-resolution structural profiling of endogenous cellular macromolecules"

**Affiliations**

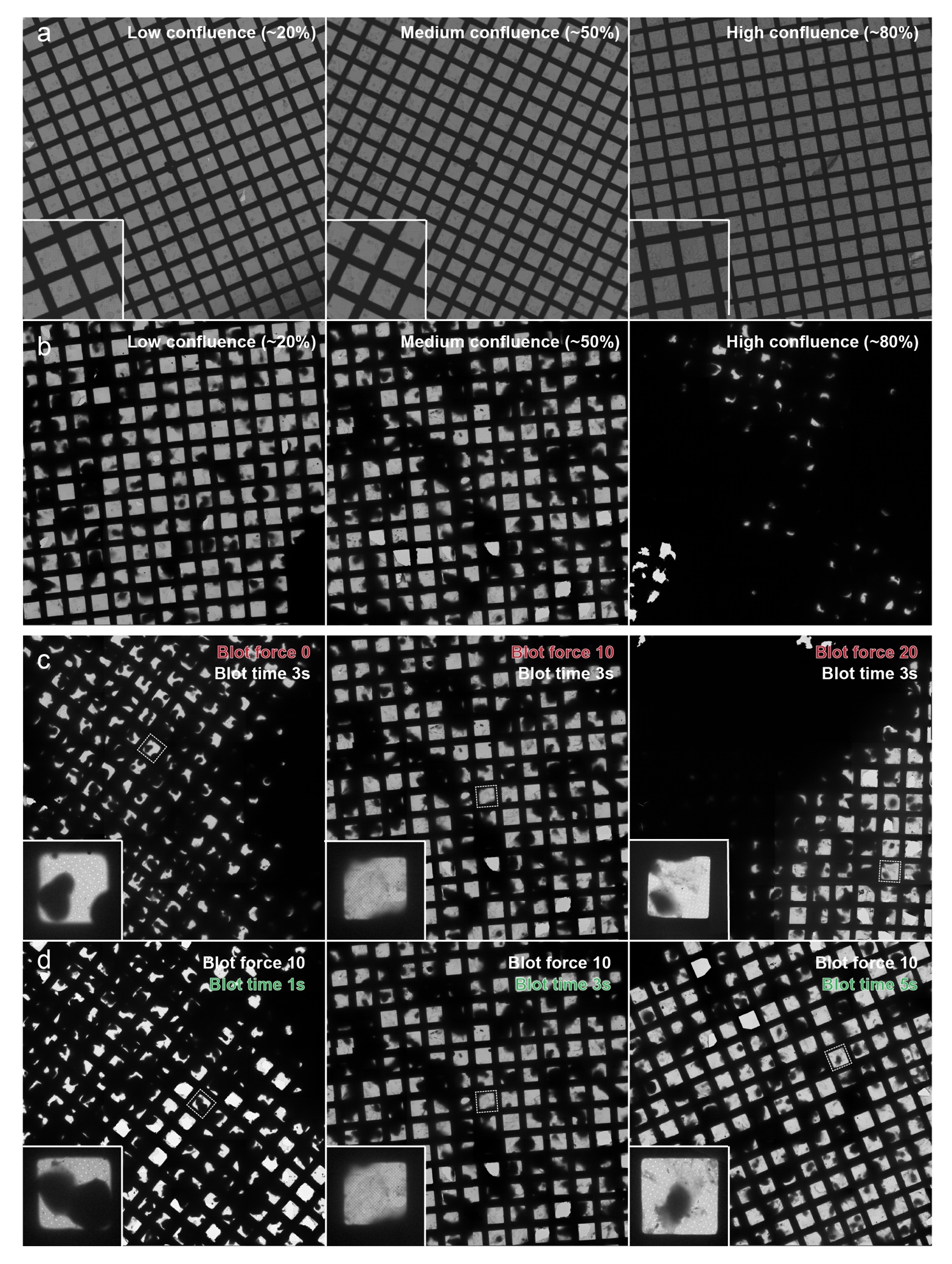

**Supplementary Fig. 1 | Optimization of cryoGO sample preparation. a,** Light microscopy images of cells attached on EM grids at low (~20%, left), medium (~50%, middle), and high (~80%, right) confluence. Insets show zoomed-in views of cell distribution. **b**, Corresponding low-magnification cryo-EM views of the grids. **c**, Low-magnification cryo-EM views of the grids prepared with varying blot forces (blot time fixed at 3 s): blot force 0 (Left), blot force 10 (Middle, as shown in b), and blot force 20 (Right). **d**, Low-magnification cryo-EM views of the grids prepared with varying blot times (blot force fixed at 10): blot time 1 s (Left), blot time 3 s (Middle, as shown in c), and blot time 5 s (Right). Insets show magnified views of representative individual grid squares.

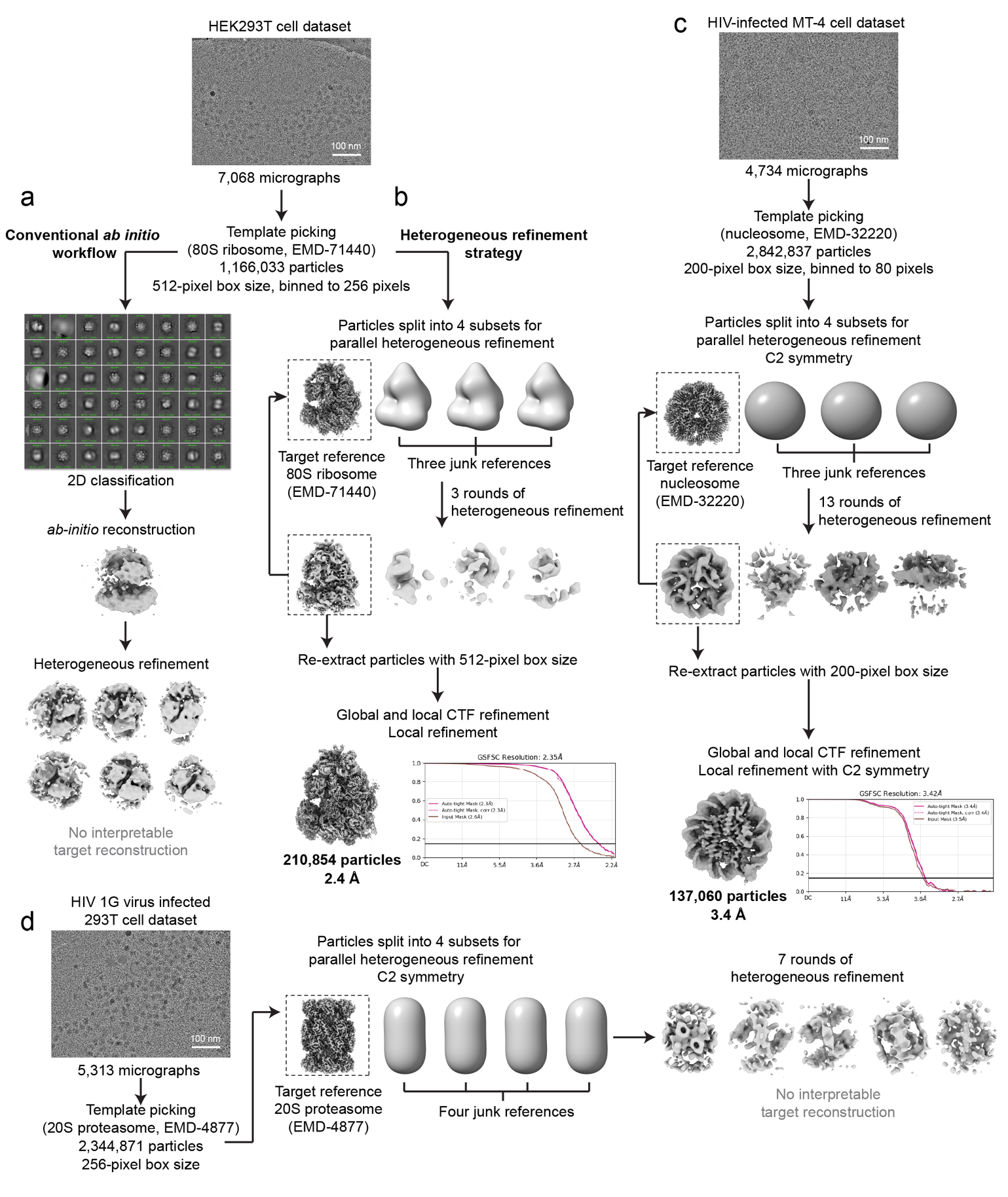

**Supplementary Fig. 2 | Evaluation of conventional *ab initio*** **and heterogeneous refinement-based processing workflows for cryoGO datasets.** **a**, Conventional *ab initio* reconstruction workflow applied to 80S ribosome particles from the HEK293T cryoGO dataset. Particles were initially detected by template picking, followed by 2D classification, *ab initio* reconstruction, and heterogeneous refinement. This workflow failed to recover an interpretable reconstruction. **b**, Application of the heterogeneous refinement strategy targeting the 80S ribosome. After 3 rounds of heterogeneous refinement against an 80S ribosome target reference and three low-pass filtered “junk” references, the retained particles were re-extracted and subjected to global and local CTF refinements, yielding a final reconstruction of the 80S ribosome at 2.3 Å resolution. **c**, Application of the heterogeneous refinement strategy targeting the nucleosome. Using a nucleosome reference and three junk references, 13 rounds of heterogeneous refinement and subsequent CTF and local refinements produced a final reconstruction at 3.4 Å resolution. **d**, Application of the heterogeneous refinement strategy targeting the proteasome. Despite employing a proteasome target reference and 7 rounds of heterogeneous refinement, the process failed to converge into an interpretable target reconstruction.

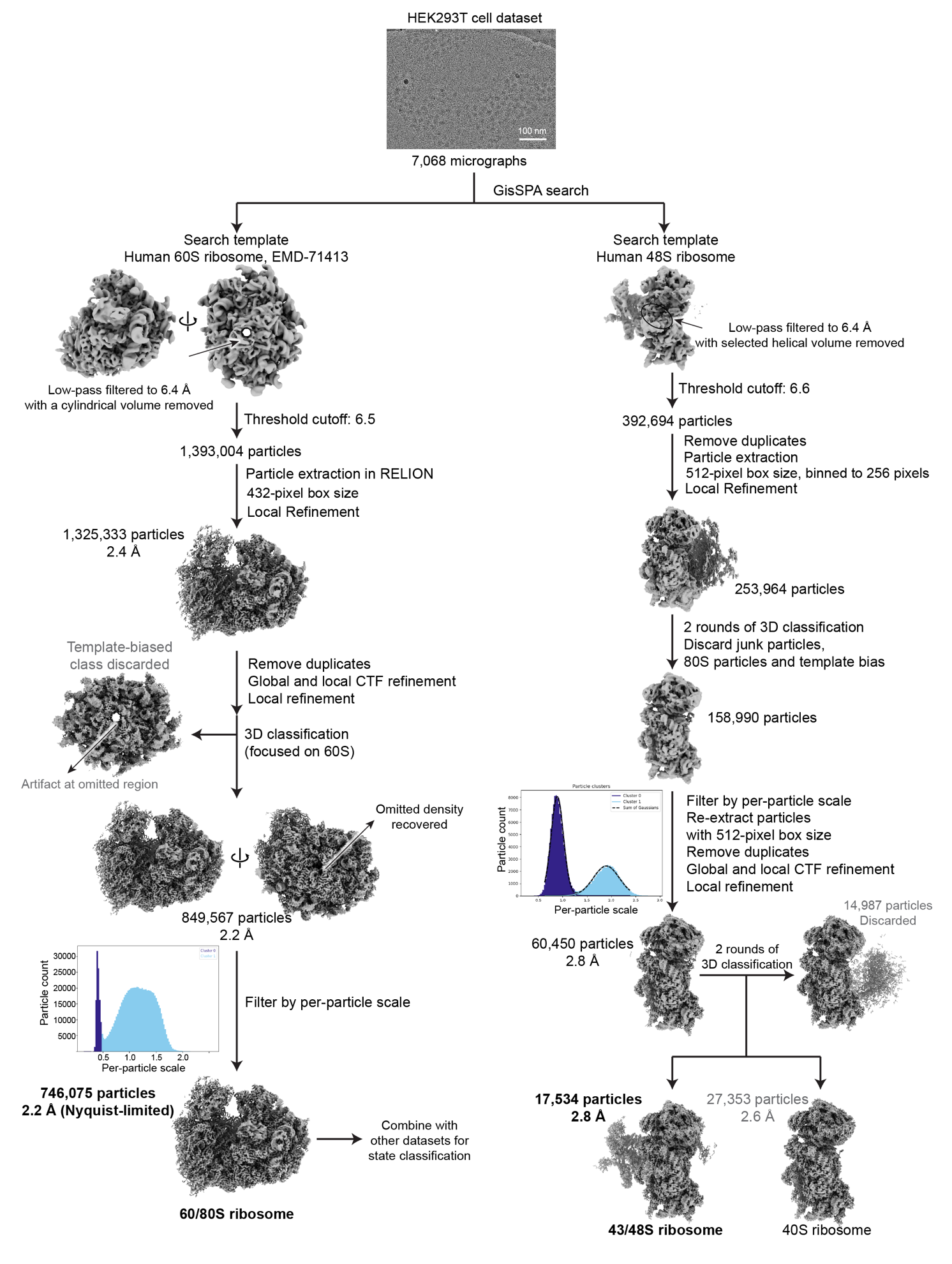

**Supplementary Fig. 3 | High-resolution 2D template-matching workflow using GisSPA for reconstructions of 60S/80S and 43S/48S ribosomes.** Left branch: Data processing pipeline for the 60S/80S ribosome. A low-pass filtered human 60S subunit reference with a cylindrical volume omitted was used as the search template with GisSPA. Subsequent 3D classification successfully separated true target particles from false positives. Following CTF refinement and per-particle scale filtering, a final 60S/80S ribosome reconstruction was achieved at a Nyquist-limited resolution of 2.2 Å. Processing workflows for 60S/80S ribosomes from all other datasets are summarized in Supplementary Fig. 9. Right branch: Data processing pipeline for the 43S/48S ribosome. The search was performed targeting translation initiation complexes using a low-pass filtered human 48S ribosome reference with an RNA duplex region omitted. Following 3D classification to separate false positives and 80S particles, the dataset was filtered by per-particle scale. After re-extraction and CTF refinements, additional rounds of 3D classification and local refinement were used to reconstruct the 43S/48S pre-initiation complex.

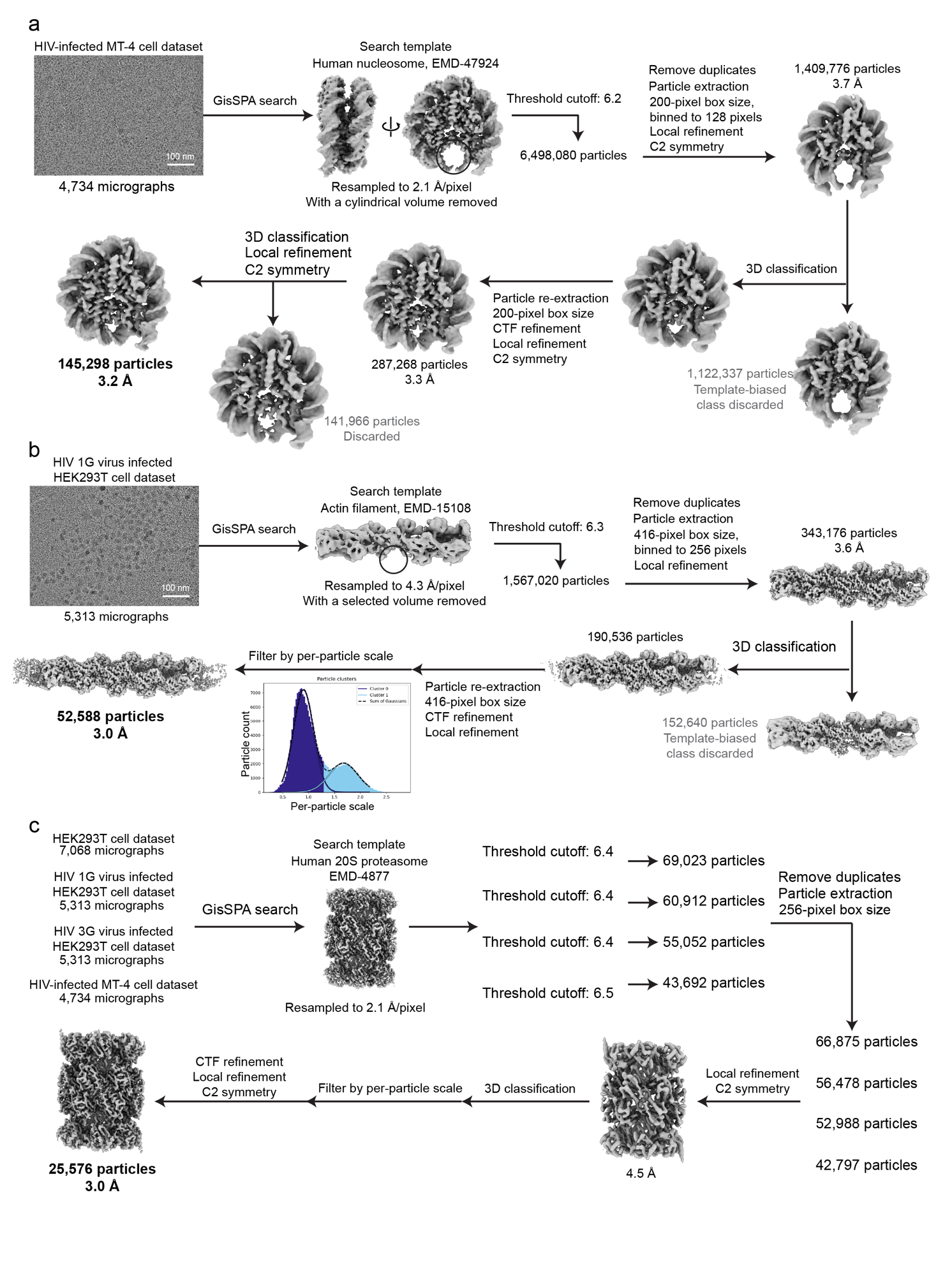

**Supplementary Fig. 4 | High-resolution 2D template-matching workflows using GisSPA for reconstructions of nucleosomes, actin filaments, and 20S proteasomes. a,** Data processing pipeline for the nucleosome. A resampled human nucleosome reference with a cylindrical volume omitted was used as the search template. Following particle extraction and local refinement, 3D classification was performed to remove template-biased particles. The retained particles were re-extracted, subjected to CTF refinement and local refinement, and further classified to yield a final nucleosome reconstruction at 3.2 Å resolution. **b,** Data processing pipeline for actin filaments. A resampled actin filament reference with selected density omitted was used as the search template. After particle extraction and local refinement, 3D classification was used to discard template-biased classes. The retained particles were re-extracted, refined, and filtered by per-particle scale, yielding a final actin filament reconstruction at 3.0 Å resolution. **c,** Data processing pipeline for 20S proteasomes. A resampled human 20S proteasome reference was used as the search template across four datasets. Particles were extracted and combined for local refinement. Followed by 3D classification and per-particle scale filtering, the retained particles were refined to 3.0 Å resolution.

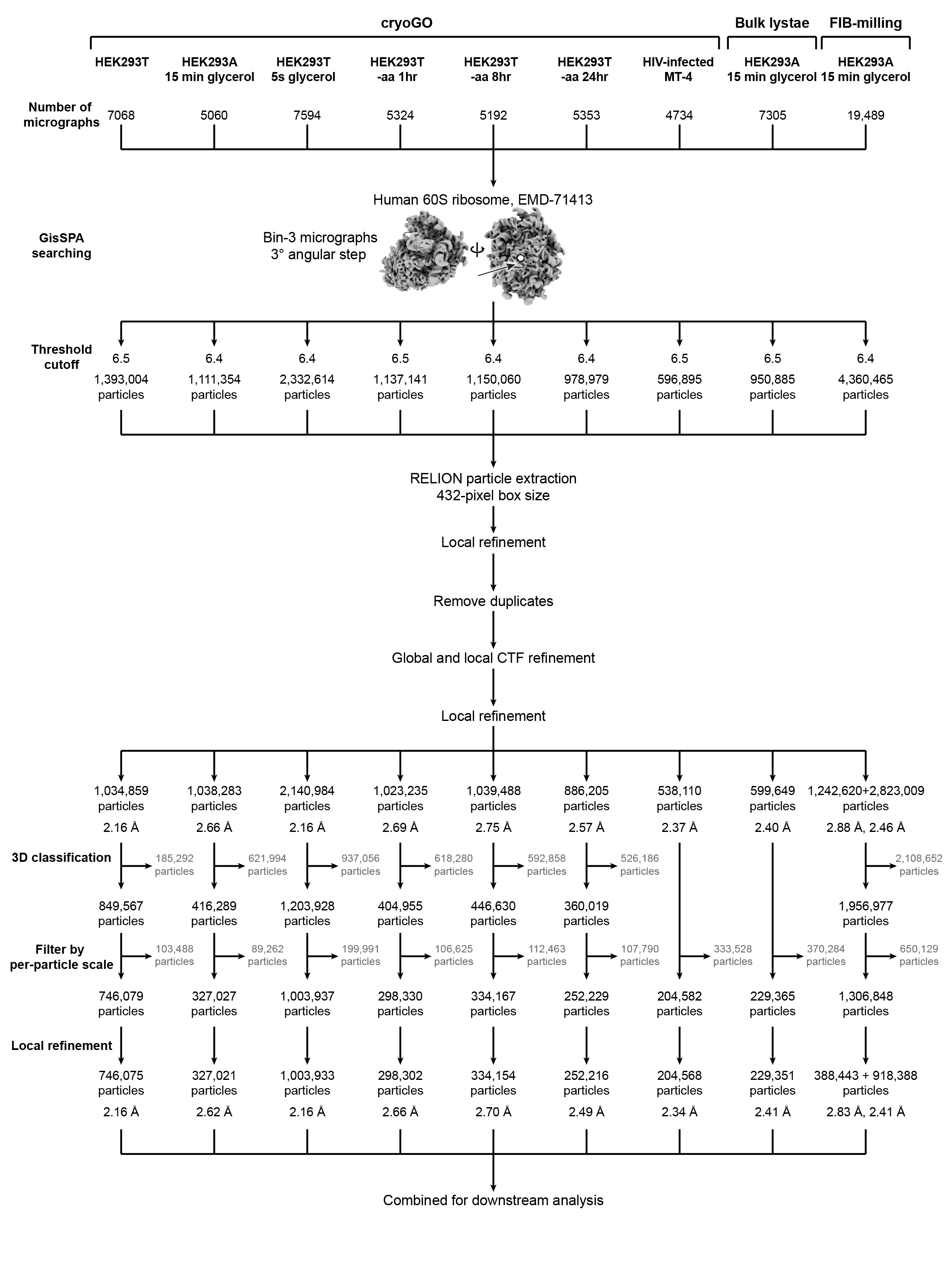

**Supplementary Fig. 5 | Data processing workflow for 60S/80S ribosomes across all evaluated datasets.**

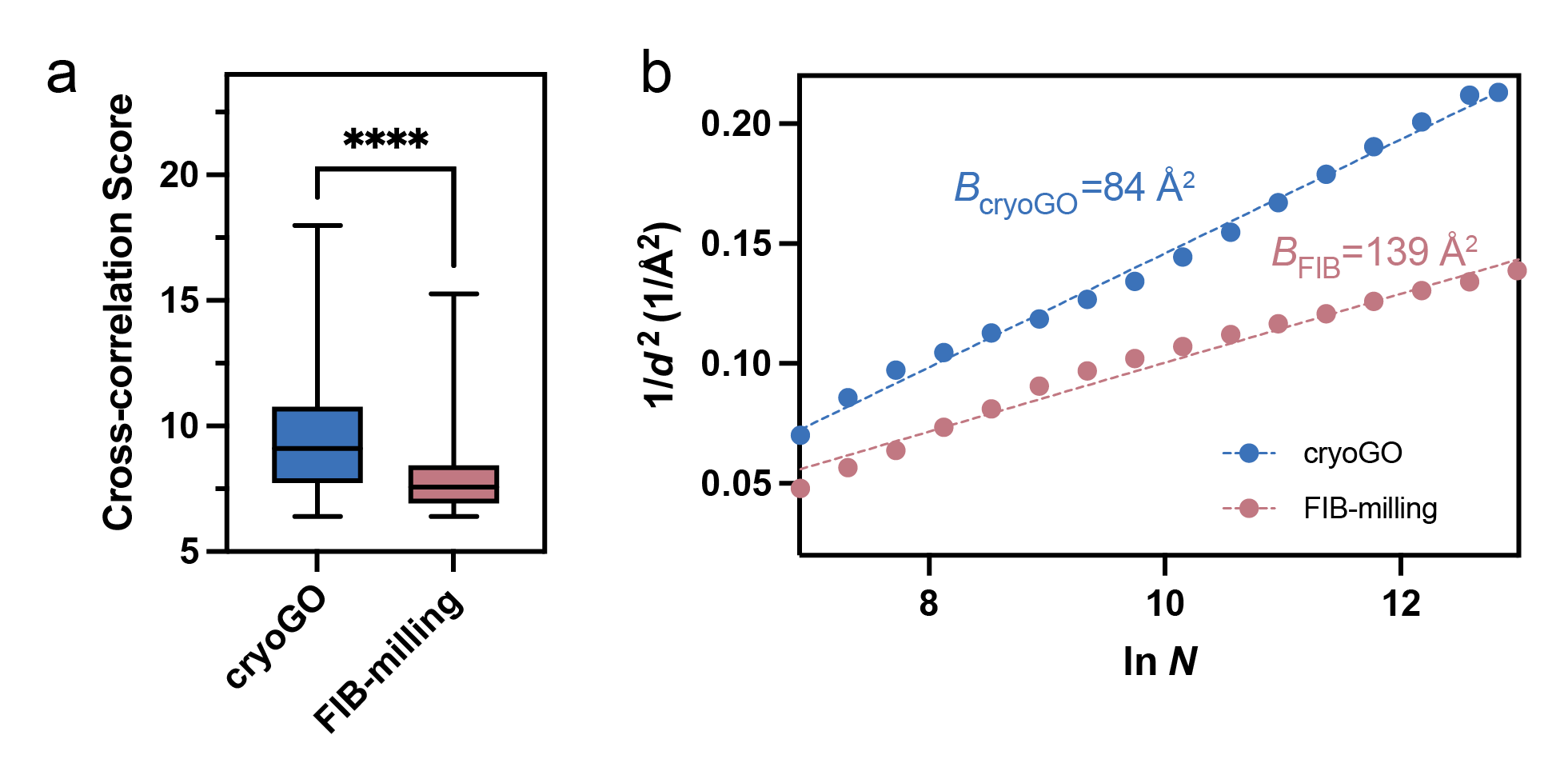

**Supplementary Fig. 6 | Quantitative comparison of data quality between cryoGO and FIB-milling samples. a**, Comparison of per-particle cross-correlation scores of ribosome particles computed during GisSPA search for cryoGO and FIB-milled lamellae datasets. Box plots indicate the median (center line), interquartile range (box limits), and extreme values (whiskers). *****p* < 0.0001. **b**, Rosenthal-Henderson plots comparing global resolution as a function of particle number for 80S ribosome reconstructions from cryoGO (blue) and FIB-milling (red) datasets. The y-axis represents the squared spatial frequency (1/*d*^2^), where 1/*d* is the spatial frequency at which the Fourier Shell Correlation (FSC) reaches 0.143. The x-axis (ln*N*) denotes the natural logarithm of the ribosome particle number used for the reconstruction. The corresponding RH B-factors (*B*) are derived from the slopes.

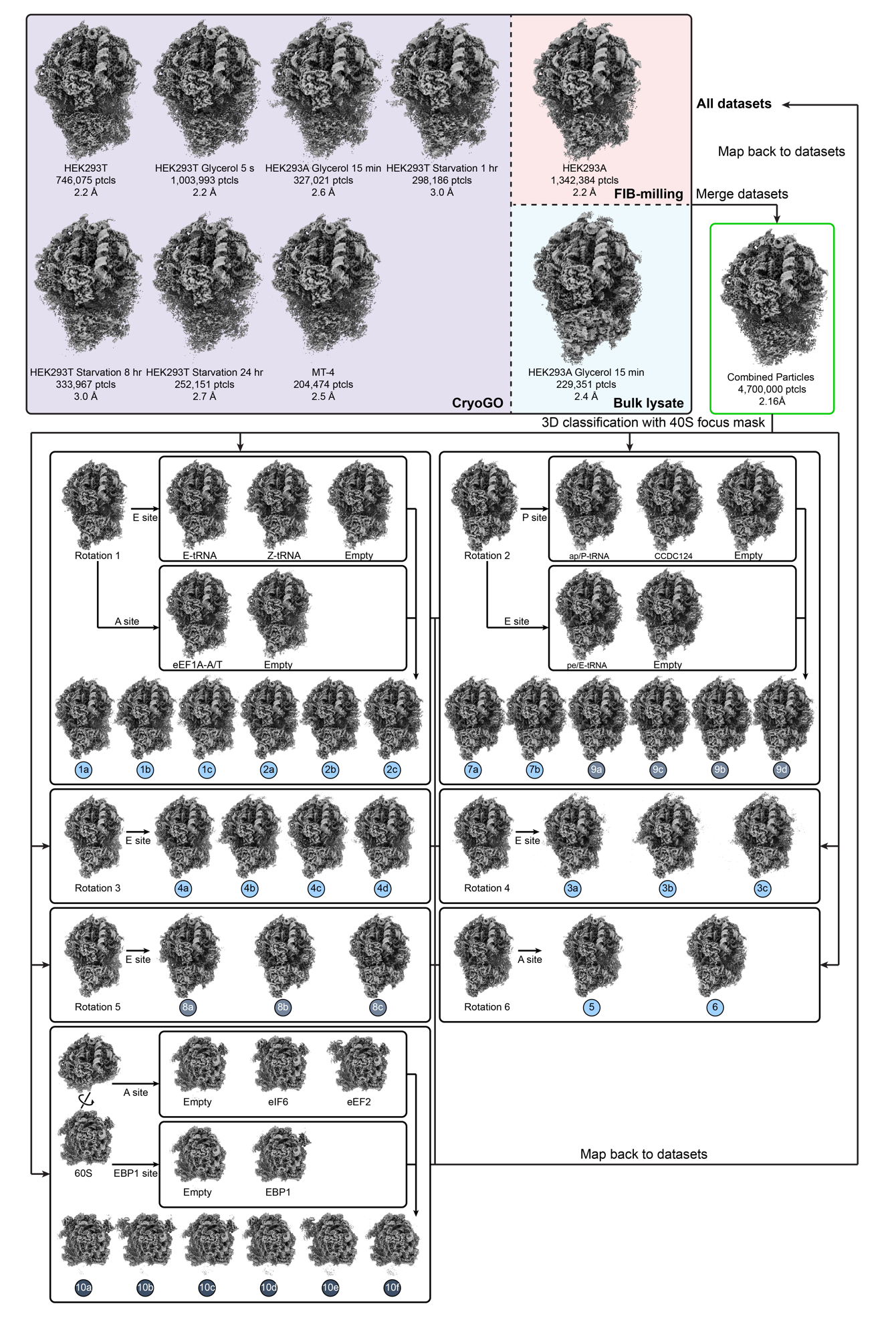

**Supplementary Fig. 7 | Hierarchical 3D classification strategy for resolving the ribosomal conformational and compositional heterogeneity.** All 80S ribosome particles from independent cryoGO datasets were pooled together and were then subjected to extensive hierarchical 3D classification using various focused masks. Following the completion of the classification pipeline, individual particles from these final classes were mapped back to their original respective datasets.

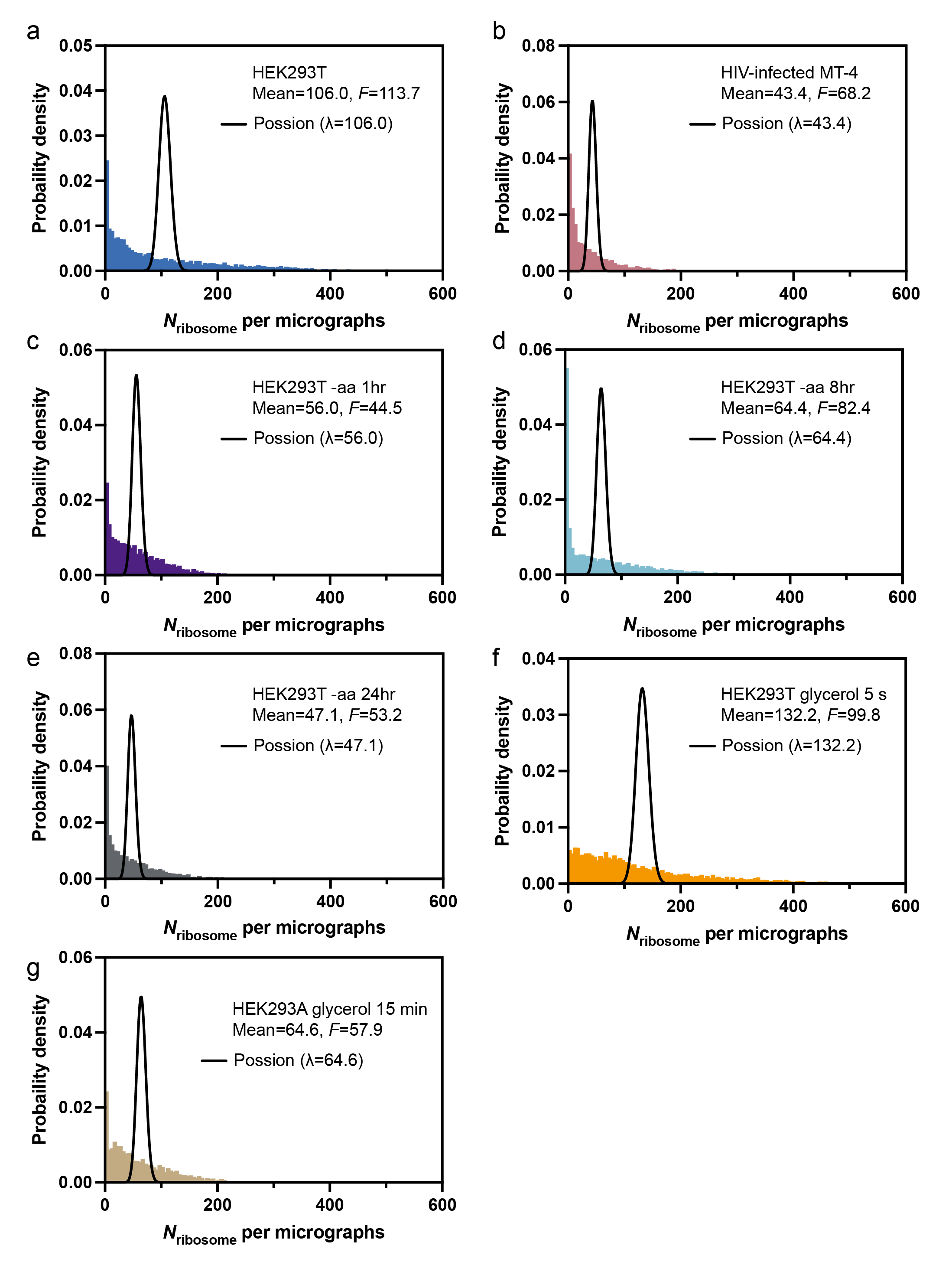

**Supplementary Fig. 8 | Per-micrograph ribosome count distributions for all evaluated cryoGO datasets. a,** HEK293T**; b,** HIV-infected MT-4**; c,** HEK293T -aa 1 hr**; d,** HEK293T -aa 8 hr**; e,** HEK293T -aa 24 hr**; f,** HEK293T glycerol 5s**; c,** HEK293A glycerol 15 min**.**

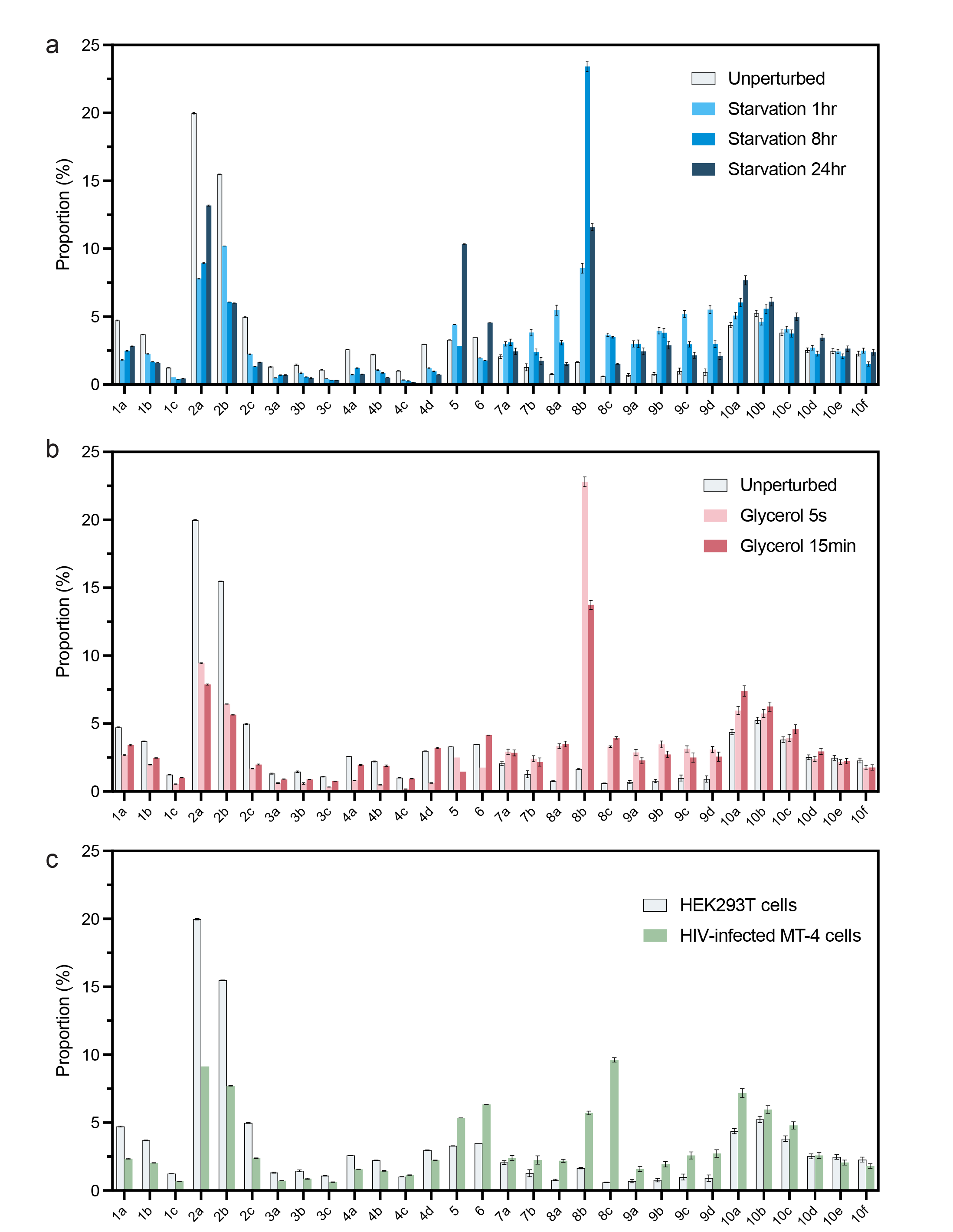

**Supplementary Fig. 9 | Quantitative profiling of ribosomal state redistributions under physiological stress and viral infection.** **a**, Bar chart showing the relative proportions of individual ribosomal states in unperturbed HEK293T cells and cells subjected to amino acid starvation for 1 h, 8 h, and 24 h. **b**, Bar chart showing the relative proportions of individual ribosomal states in unperturbed cells and cells treated with 5% glycerol (5 s and 15 min). **c**, Bar chart comparing the relative proportions of individual ribosomal states between unperturbed HEK293T cells and HIV-infected MT-4 cells. For all panels, data are presented as mean ± standard deviation derived from three independent classification analyses.

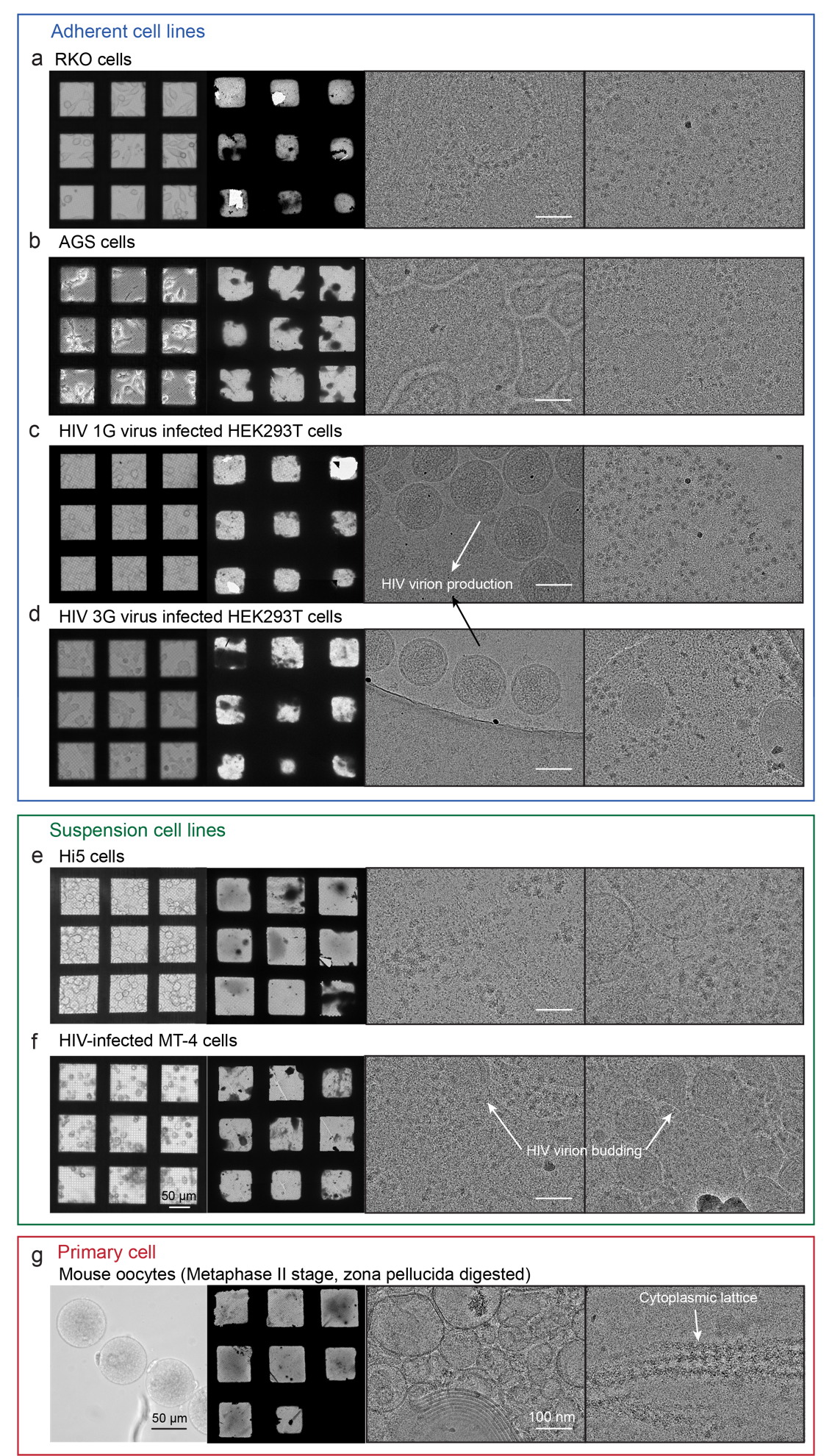

**Supplementary Fig. 10 | Broad applicability of cryoGO across diverse adherent, suspension, and primary cell samples.** **a–g**, Representative images demonstrating the successful application of the cryoGO workflow to a wide array of cell types, including adherent cell lines (**a**, human colon carcinoma RKO cells; **b**, human gastric adenocarcinoma AGS cells; **c**, HIV-1G virus infected HEK293T cells; **d**, HIV-3G virus infected HEK293T cells), suspension cell lines (**e**, insect ovary Hi5 cells; **f**, HIV-infected human T-cell leukemia MT-4 cells), and a primary cell model (**g**, mouse Metaphase II oocytes). For all panels, images from left to right: bright-field light microscopy of cells; low-magnification cryo-EM montages; and two representative high-magnification cryo-EM micrographs. The micrographs capture distinct cell-specific features, such as HIV virion production and budding events in the infected cells (arrows in **c**, **d**, and **f**) and the cytoplasmic lattice unique to the mammalian oocyte^45^ (arrow in **g**).

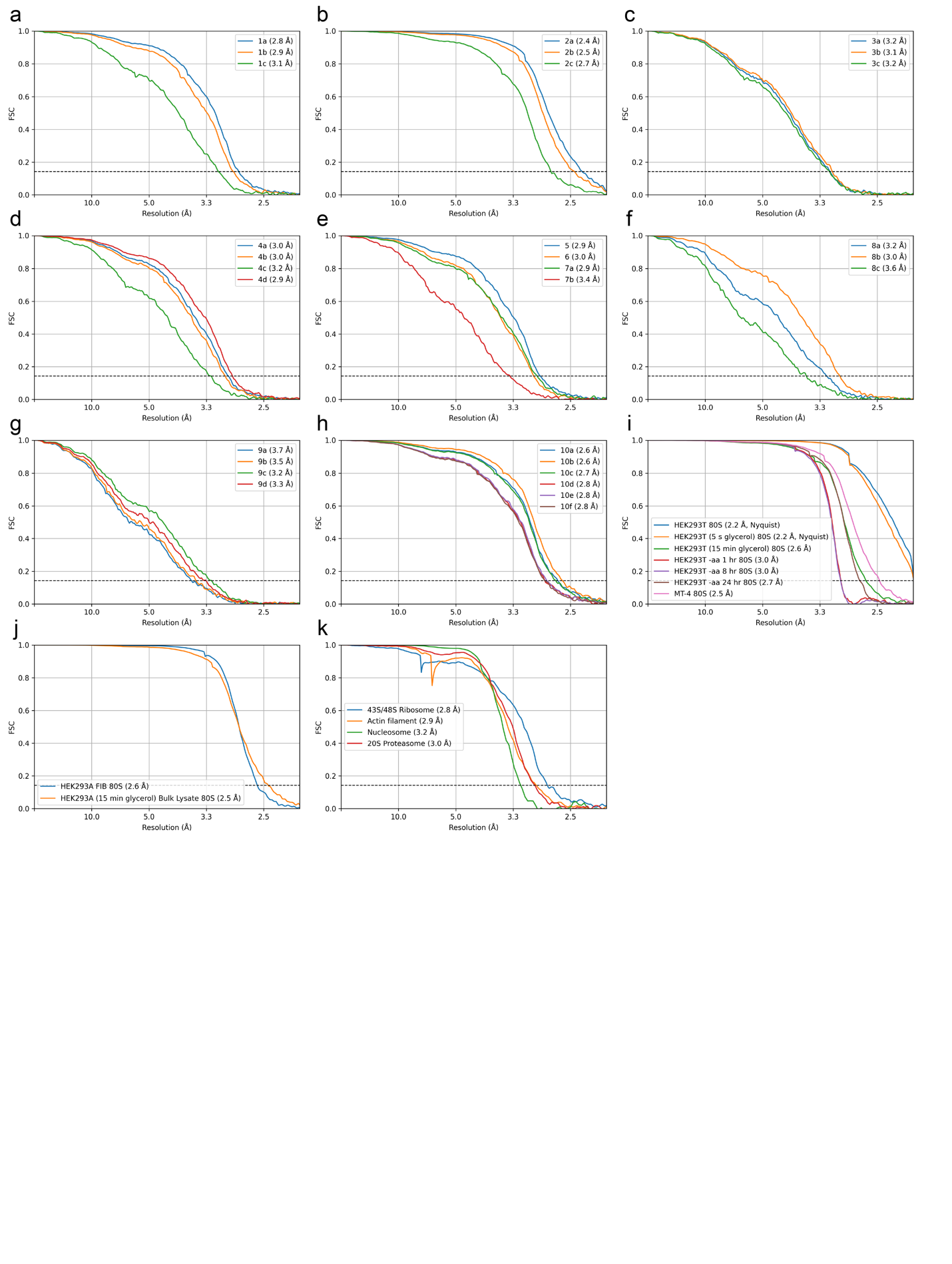

**Supplementary Fig. 11 | Fourier shell correlation (FSC) curves for all cryo-EM reconstructions. a,** Ribosome states 1a–1c; **b,** Ribosome states 2a–2c; **c,** Ribosome states 3a–3c; **d,** Ribosome states 4a–4d; **e,** Ribosome states 5, 6, 7a and 7b; **f,** Ribosome states 8a–8c; **g,** Ribosome states 9a–9d; **h,** Ribosome states 10a–10f; **i,** consensus 80S ribosome reconstructions across the diverse cryoGO datasets; **j,** consensus 80S ribosome reconstructions from FIB-milled lamellae and bulk lysate preparations; **k,** aditional endogenous macromolecular targets resolved using the cryoGO workflow, including the 43S/48S ribosome, actin filament, nucleosome, and 20S proteasome. For all panels, the gold-standard FSC=0.143 criterion is indicated by the horizontal dashed black lines.

**Supplementary Table 1 | Summary of particle yields and reconstruction resolutions using heterogeneous refinement and high-resolution template-matching processing strategies.**

| **Processing strategy** | **80S ribosome** | | **Nucleosome** | | **20S proteasome** | |
| --- | --- | --- | --- | --- | --- | --- |
|  | **Particle number** | **Resolution (Å)** | **Particle number** | **Resolution (Å)** | **Particle number** | **Resolution (Å)** |
| **Heterogeneous refinement** | 210,854 | 2.4 | 137,060 | 3.4 | No interpretable target reconstruction | |
| **High-resolution template matching** | 746,075 | 2.2 (Nyquist-limited ) | 145,298 | 3.2 | 25,576 | 3.0 |

**Supplementary Table 2 | Summary statistics of ribosome per-particle cross-correlation scores computed from GisSPA.**

| **Sample** | **Particle number** | **Mean** | **Standard deviation** | **Median** | **Minimum** | **Maximum** |
| --- | --- | --- | --- | --- | --- | --- |
| **cryoGO** | 746,075 | 9.4 | 2.0 | 9.1 | 6.4 | 18.0 |
| **FIB-milling** | 388,443 | 7.8 | 1.1 | 7.6 | 6.4 | 15.3 |

**Supplementary Table 3 | Percentage distributions of the 60S and 80S ribosomal states**

| **States** | | | **States distribution (%)** | | | | | | | | | | | |
| --- | --- | --- | --- | --- | --- | --- | --- | --- | --- | --- | --- | --- | --- | --- |
|  |  |  | **cryoGO**  **HEK293T** | | | **cryoGO**  **HEK293A glycerol 15 min** | | | **FIB-milling HEK293A glycerol 15 min** | | | **Bulk lysate**  **HEK293A glycerol 15 min** | | |
| **Translating 80S** | 1a | P-E | 4.70 | 4.76 | 4.71 | 3.37 | 3.48 | 3.42 | 3.98 | 4.09 | 4.02 | 3.93 | 4.06 | 3.97 |
|  | 1b | P-Z | 3.71 | 3.72 | 3.67 | 2.45 | 2.49 | 2.46 | 2.24 | 2.26 | 2.23 | 0.52 | 0.54 | 0.53 |
|  | 1c | P | 1.23 | 1.26 | 1.24 | 1.00 | 1.04 | 1.02 | 0.68 | 0.71 | 0.69 | 0.42 | 0.41 | 0.40 |
|  | 2a | eEF1A-AT-P-E-Sampling | 19.94 | 20.03 | 19.96 | 7.83 | 7.91 | 7.89 | 12.78 | 12.83 | 12.81 | 6.21 | 6.23 | 6.21 |
|  | 2b | eEF1A-AT-P-Z-Sampling | 15.50 | 15.49 | 15.45 | 5.69 | 5.64 | 5.64 | 7.39 | 7.37 | 7.35 | 1.08 | 1.08 | 1.08 |
|  | 2c | eEF1A-AT-P-Sampling | 4.95 | 5.02 | 4.98 | 1.95 | 2.01 | 2.00 | 2.14 | 2.18 | 2.15 | 0.80 | 0.80 | 0.79 |
|  | 3a | eEF1A-AT-P-E-Decoding | 1.33 | 1.27 | 1.35 | 0.91 | 0.85 | 0.90 | 1.21 | 1.13 | 1.20 | 1.97 | 1.89 | 2.01 |
|  | 3b | eEF1A-AT-P-Z-Decoding | 1.45 | 1.39 | 1.52 | 0.88 | 0.85 | 0.90 | 0.86 | 0.82 | 0.91 | 0.23 | 0.23 | 0.17 |
|  | 3c | eEF1A-AT-P-Decoding | 1.10 | 1.06 | 1.11 | 0.77 | 0.75 | 0.75 | 0.66 | 0.63 | 0.65 | 0.37 | 0.36 | 0.38 |
|  | 4a | A-P-E | 2.57 | 2.60 | 2.57 | 1.93 | 2.01 | 1.93 | 2.27 | 2.32 | 2.26 | 1.76 | 1.78 | 1.75 |
|  | 4b | A-P-Z | 2.20 | 2.25 | 2.20 | 1.87 | 1.96 | 1.86 | 1.43 | 1.48 | 1.42 | 0.25 | 0.26 | 0.25 |
|  | 4c | A-P-eIF5A | 1.02 | 1.03 | 1.00 | 0.94 | 0.97 | 0.92 | 0.68 | 0.70 | 0.66 | 0.86 | 0.91 | 0.87 |
|  | 4d | A-P | 2.97 | 2.99 | 2.97 | 3.17 | 3.27 | 3.19 | 2.05 | 2.11 | 2.05 | 0.55 | 0.58 | 0.56 |
|  | 5 | AP-PE | 3.29 | 3.30 | 3.29 | 1.45 | 1.45 | 1.45 | 1.64 | 1.65 | 1.63 | 1.11 | 1.12 | 1.11 |
|  | 6 | eEF2-AP-PE | 3.48 | 3.48 | 3.49 | 4.15 | 4.15 | 4.16 | 6.23 | 6.25 | 6.24 | 0.32 | 0.31 | 0.32 |
|  | 7a | eEF2-apP-peE | 1.96 | 2.22 | 2.00 | 2.65 | 3.07 | 2.80 | 2.91 | 3.29 | 3.03 | 1.65 | 1.76 | 1.68 |
|  | 7b | eEF2-apP | 1.10 | 1.57 | 1.16 | 1.93 | 2.52 | 2.06 | 1.74 | 2.25 | 1.84 | 1.63 | 1.84 | 1.80 |
| **Non-translating 80S** | 8a | Hiber 1: eEF2 | 0.83 | 0.76 | 0.75 | 3.74 | 3.42 | 3.33 | 2.15 | 1.99 | 1.98 | 1.16 | 1.09 | 1.08 |
|  | 8b | Hiber 1: eEF2-E | 1.70 | 1.64 | 1.60 | 14.08 | 13.72 | 13.42 | 15.85 | 15.51 | 15.37 | 8.36 | 8.32 | 8.22 |
|  | 8c | Hiber 1: eEF2-eIF5A | 0.63 | 0.60 | 0.59 | 4.05 | 3.92 | 3.88 | 3.71 | 3.63 | 3.61 | 47.58 | 47.37 | 46.90 |
|  | 9a | Hiber 2: eEF2-CCDC124-E | 0.62 | 0.84 | 0.63 | 2.06 | 2.54 | 2.25 | 1.97 | 2.36 | 2.08 | 2.13 | 2.34 | 2.15 |
|  | 9b | Hiber 2: eEF2-E | 0.68 | 0.92 | 0.71 | 2.48 | 2.98 | 2.74 | 2.38 | 2.85 | 2.57 | 1.79 | 1.98 | 1.86 |
|  | 9c | Hiber 2: eEF2-CCDC124 | 0.85 | 1.25 | 0.88 | 2.21 | 2.87 | 2.42 | 1.70 | 2.17 | 1.78 | 2.86 | 3.09 | 3.12 |
|  | 9d | Hiber 2: eEF2 | 0.77 | 1.18 | 0.80 | 2.27 | 2.93 | 2.49 | 1.72 | 2.25 | 1.85 | 2.47 | 2.72 | 2.81 |
| **60S** | 10a | 60S-EBP1-eIF6 | 4.49 | 4.15 | 4.49 | 7.63 | 6.95 | 7.61 | 6.30 | 5.67 | 6.30 | 2.51 | 2.30 | 2.51 |
|  | 10b | 60S-EBP1-eEF2 | 5.39 | 4.97 | 5.36 | 3.37 | 3.48 | 3.42 | 3.98 | 4.09 | 4.02 | 3.93 | 4.06 | 3.97 |
|  | 10c | 60S-EBP1 | 3.94 | 3.58 | 3.94 | 2.45 | 2.49 | 2.46 | 2.24 | 2.26 | 2.23 | 0.52 | 0.54 | 0.53 |
|  | 10d | 60S-eIF6 | 2.63 | 2.32 | 2.62 | 1.00 | 1.04 | 1.02 | 0.68 | 0.71 | 0.69 | 0.42 | 0.41 | 0.40 |
|  | 10e | 60S-eEF2 | 2.58 | 2.27 | 2.57 | 7.83 | 7.91 | 7.89 | 12.78 | 12.83 | 12.81 | 6.21 | 6.23 | 6.21 |
|  | 10f | 60S | 2.39 | 2.07 | 2.38 | 5.69 | 5.64 | 5.64 | 7.39 | 7.37 | 7.35 | 1.08 | 1.08 | 1.08 |

**Supplementary Table 3 continued**

| **States** | | **States distribution (%)** | | | | | | | | | | | | | | |
| --- | --- | --- | --- | --- | --- | --- | --- | --- | --- | --- | --- | --- | --- | --- | --- | --- |
|  |  | **HEK293T starvation 1hr** | | | **HEK293T starvation 1hr** | | | **HEK293T starvation 1hr** | | | **HEK293T glycerol 5s** | | | **HIV-infected MT-4 cell** | | |
| **Translating 80S** | 1a | 1.80 | 1.84 | 1.82 | 2.46 | 2.51 | 2.48 | 2.79 | 2.84 | 2.82 | 2.65 | 2.71 | 2.68 | 2.31 | 2.38 | 2.35 |
|  | 1b | 2.26 | 2.28 | 2.25 | 1.67 | 1.69 | 1.67 | 1.60 | 1.61 | 1.60 | 1.96 | 1.99 | 1.96 | 2.03 | 2.05 | 2.03 |
|  | 1c | 0.53 | 0.54 | 0.54 | 0.40 | 0.41 | 0.40 | 0.44 | 0.46 | 0.45 | 0.54 | 0.56 | 0.55 | 0.67 | 0.70 | 0.68 |
|  | 2a | 7.78 | 7.83 | 7.82 | 8.89 | 8.96 | 8.97 | 13.14 | 13.22 | 13.19 | 9.41 | 9.48 | 9.44 | 9.13 | 9.13 | 9.13 |
|  | 2b | 10.21 | 10.21 | 10.19 | 6.08 | 6.07 | 6.07 | 6.02 | 6.00 | 5.99 | 6.46 | 6.45 | 6.43 | 7.70 | 7.74 | 7.70 |
|  | 2c | 2.20 | 2.25 | 2.24 | 1.33 | 1.35 | 1.34 | 1.64 | 1.65 | 1.63 | 1.67 | 1.70 | 1.69 | 2.37 | 2.41 | 2.40 |
|  | 3a | 0.51 | 0.48 | 0.50 | 0.72 | 0.68 | 0.71 | 0.72 | 0.68 | 0.73 | 0.63 | 0.59 | 0.64 | 0.74 | 0.71 | 0.73 |
|  | 3b | 0.87 | 0.82 | 0.93 | 0.58 | 0.55 | 0.57 | 0.47 | 0.44 | 0.55 | 0.57 | 0.54 | 0.65 | 0.89 | 0.83 | 0.89 |
|  | 3c | 0.43 | 0.41 | 0.43 | 0.34 | 0.33 | 0.34 | 0.34 | 0.31 | 0.33 | 0.34 | 0.32 | 0.34 | 0.64 | 0.60 | 0.63 |
|  | 4a | 0.72 | 0.76 | 0.72 | 1.21 | 1.25 | 1.21 | 0.76 | 0.78 | 0.76 | 0.80 | 0.83 | 0.80 | 1.56 | 1.57 | 1.56 |
|  | 4b | 1.05 | 1.10 | 1.05 | 0.85 | 0.88 | 0.85 | 0.51 | 0.53 | 0.51 | 0.49 | 0.52 | 0.48 | 1.44 | 1.48 | 1.43 |
|  | 4c | 0.33 | 0.36 | 0.33 | 0.27 | 0.29 | 0.27 | 0.18 | 0.18 | 0.17 | 0.17 | 0.19 | 0.17 | 1.14 | 1.17 | 1.13 |
|  | 4d | 1.18 | 1.25 | 1.18 | 0.96 | 1.00 | 0.95 | 0.72 | 0.73 | 0.73 | 0.61 | 0.64 | 0.61 | 2.22 | 2.24 | 2.24 |
|  | 5 | 4.41 | 4.42 | 4.41 | 2.85 | 2.85 | 2.84 | 10.36 | 10.36 | 10.33 | 2.50 | 2.50 | 2.50 | 5.35 | 5.34 | 5.34 |
|  | 6 | 1.96 | 1.96 | 1.97 | 1.77 | 1.77 | 1.78 | 4.55 | 4.55 | 4.56 | 1.76 | 1.76 | 1.76 | 6.33 | 6.32 | 6.34 |
|  | 7a | 2.84 | 3.18 | 2.98 | 2.89 | 3.37 | 3.08 | 2.26 | 2.71 | 2.37 | 2.75 | 3.12 | 2.90 | 2.28 | 2.60 | 2.32 |
|  | 7b | 3.59 | 4.06 | 3.83 | 2.22 | 2.64 | 2.34 | 1.57 | 2.03 | 1.65 | 2.23 | 2.66 | 2.35 | 2.00 | 2.59 | 2.15 |
| **Non-translating 80S** | 8a | 5.88 | 5.41 | 5.14 | 3.27 | 3.06 | 2.95 | 1.61 | 1.52 | 1.41 | 3.52 | 3.33 | 3.17 | 2.30 | 2.20 | 2.07 |
|  | 8b | 8.92 | 8.56 | 8.21 | 23.78 | 23.40 | 23.05 | 11.78 | 11.71 | 11.30 | 23.11 | 22.90 | 22.41 | 5.84 | 5.73 | 5.57 |
|  | 8c | 3.80 | 3.64 | 3.55 | 3.55 | 3.49 | 3.43 | 1.57 | 1.54 | 1.49 | 3.35 | 3.31 | 3.23 | 9.78 | 9.64 | 9.45 |
|  | 9a | 2.78 | 3.24 | 2.97 | 2.75 | 3.31 | 2.94 | 2.24 | 2.73 | 2.37 | 2.66 | 3.11 | 2.83 | 1.44 | 1.79 | 1.51 |
|  | 9b | 3.72 | 4.19 | 3.99 | 3.51 | 4.15 | 3.79 | 2.65 | 3.19 | 2.85 | 3.21 | 3.71 | 3.48 | 1.77 | 2.16 | 1.88 |
|  | 9c | 4.91 | 5.43 | 5.24 | 2.78 | 3.18 | 2.94 | 1.99 | 2.41 | 2.07 | 2.93 | 3.39 | 3.09 | 2.36 | 2.86 | 2.54 |
|  | 9d | 5.19 | 5.74 | 5.61 | 2.75 | 3.24 | 2.97 | 1.90 | 2.37 | 2.01 | 2.87 | 3.32 | 3.05 | 2.46 | 3.01 | 2.72 |
| **60S** | 10a | 5.22 | 4.80 | 5.21 | 6.22 | 5.67 | 6.23 | 7.88 | 7.29 | 7.87 | 6.13 | 5.61 | 6.12 | 7.36 | 6.80 | 7.38 |
|  | 10b | 4.77 | 4.35 | 4.74 | 5.79 | 5.19 | 5.77 | 6.31 | 5.75 | 6.28 | 5.93 | 5.39 | 5.90 | 6.15 | 5.65 | 6.11 |
|  | 10c | 4.20 | 3.83 | 4.21 | 3.91 | 3.46 | 3.92 | 5.14 | 4.66 | 5.16 | 4.09 | 3.63 | 4.10 | 4.95 | 4.49 | 4.95 |
|  | 10d | 2.80 | 2.51 | 2.82 | 2.39 | 2.07 | 2.38 | 3.59 | 3.19 | 3.59 | 2.51 | 2.19 | 2.51 | 2.71 | 2.34 | 2.70 |
|  | 10e | 2.54 | 2.25 | 2.52 | 2.19 | 1.87 | 2.18 | 2.77 | 2.41 | 2.75 | 2.28 | 1.96 | 2.26 | 2.18 | 1.85 | 2.17 |
|  | 10f | 2.61 | 2.28 | 2.61 | 1.61 | 1.33 | 1.61 | 2.50 | 2.14 | 2.51 | 1.85 | 1.57 | 1.86 | 1.90 | 1.62 | 1.91 |

**Supplementary Table 4 | Statistical analyses comparing ribosome state distributions to the FIB-milling baseline.**

|  | **Linear regression** | | | **JS divergence** | |
| --- | --- | --- | --- | --- | --- |
| **Sample** | ***R*^2^** | ***P* value** | **Slope (95% confidence interval)** | **Mean** | **Standard deviation** |
| **cryoGO** | 0.88 | <0.0001 | 0.74 (0.68 to 0.80) | 0.011 | 0.001 |
| **Bulk lysate** | 0.03 | 0.0853 | 0.45 (-0.06 to 0.97) | 0.177 | 0.001 |

**Supplementary Table 5 | Summary statistics of per-micrograph ribosome distribution**

| **Sample** | ***n* (mics)** | **⟨*N*⟩** | ***F*** | **CV²[λ]** |
| --- | --- | --- | --- | --- |
| **Bulk lysate** | 7,305 | 31.4 | 16.8 | 0.50 |
| **cryoGO** | 7,036 | 106.0 | 113.7 | 1.06 |
| **FIB-milling** | 12,492 | 73.5 | 152.1 | 2.06 |

*n* (mics), number of micrographs. ⟨*N*⟩, mean particles per micrograph. *F* = Var[*N*]/⟨*N*⟩, the Fano factor (variance-to-mean ratio). CV²[λ] = (*F*−1)/⟨*N*⟩ is the relative variance of the underlying particle density λ after removing Poisson sampling noise (compound-Poisson decomposition; CV²[λ] = 0 for a perfectly well-mixed sample) and is invariant to the absolute mean.

**Supplementary Table 6 | Cryo-EM data collection and refinement statistics for consensus maps**

|  | 80S ribosome from HEK293T  prepared by cryoGO | 80S ribosome from HEK293T glycerol 5 s  prepared by cryoGO | 80S ribosome from HEK293A glycerol 15 min  prepared by cryoGO | 80S ribosome from HEK293T -aa 1hr  prepared by cryoGO | 80S ribosome from HEK293T -aa 8hr  prepared by cryoGO | 80S ribosome from HEK293T -aa 24hr  prepared by cryoGO |
| --- | --- | --- | --- | --- | --- | --- |
|  | EMD-77427 | EMD-77458 | EMD-77459 | EMD-77462 | EMD-77463 | EMD-77464 |
| **Data collection and processing** |  |  |  |  |  |  |
| Magnification | 81,000 | 81,000 | 81,000 | 45,000 | 45,000 | 45,000 |
| Voltage (kV) | 300 | 300 | 300 | 200 | 200 | 200 |
| Electron exposure (e-/ Å^2^) | 50 | 50 | 50 | 50 | 50 | 50 |
| Defocus range (μm) | -0.5 to -1.5 | -0.5 to -1.5 | -0.5 to -1.5 | -0.5 to -1.5 | -0.5 to -1.5 | -0.5 to -1.5 |
| Pixel size (Å) | 1.068 | 1.068 | 1.068 | 0.86 | 0.86 | 0.86 |
| Micrographs (no.) | 7,036 | 7,506 | 5,047 | 5,319 | 5,176 | 4,891 |
| Final particle images (no.) | 746,024 | 1,003,882 | 327,005 | 298,174 | 333,962 | 252,141 |
| Symmetry imposed | C1 | C1 | C1 | C1 | C1 | C1 |
| Map resolution (Å) | 2.16 | 2.16 | 2.63 | 2.97 | 2.98 | 2.71 |
| FSC threshold | Nyquist | Nyquist | 0.143 | 0.143 | 0.143 | 0.143 |

**Supplementary Table 6 | Cryo-EM data collection and refinement statistics for 80S ribosome consensus maps**

|  | 80S ribosome from HIV-infected MT-4  prepared by cryoGO | 80S ribosome from HEK293A with 15 min glycerol treatment  prepared by FIB-milling | 80S ribosome from HEK293A with 15 min glycerol treatment  prepared by bulk lysate | 43S/48S ribosomes prepared by cryoGO | Nucleosome prepared by cryoGO | 20S proteasome prepared by cryoGO | Actin filament prepared by cryoGO |
| --- | --- | --- | --- | --- | --- | --- | --- |
|  | EMD-77465 | EMD-77460 | EMD-77461 | EMD-77466 | EMD-77467 | EMD-77468 | EMD-77469 |
| **Data collection and processing** |  |  |  |  |  |  |  |
| Magnification | 81,000 | 81,000 | 81,000 | 81,000 | 81,000 | 81,000 | 81,000 |
| Voltage (kV) | 300 | 300 | 300 | 300 | 300 | 300 | 300 |
| Electron exposure (e-/ Å^2^) | 50 | 50 | 50 | 50 | 50 | 50 | 50 |
| Defocus range (μm) | -0.5 to -1.5 | -0.5 to -1.5 | -0.5 to -1.5 | -0.5 to -1.5 | -0.5 to -1.5 | -0.5 to -1.5 | -0.5 to -1.5 |
| Pixel size (Å) | 1.068 | 1.068 | 1.068 | 1.068 | 1.068 | 1.068 | 1.068 |
| Micrographs (no.) | 4,717 | 18,581 | 7,208 | 7,036 | 4,717 | 10,655 | 5,264 |
| Final particle images (no.) | 204,463 | 1,306,414 | 229,338 | 17,534 | 145,298 | 25,576 | 52,588 |
| Symmetry imposed | C1 | C1 | C1 | C1 | C2 | C2 | C2 |
| Map resolution (Å) | 2.48 | 2.57 | 2.45 | 2.78 | 3.19 | 2.96 | 2.94 |
| FSC threshold | 0.143 | 0.143 | 0.143 | 0.143 | 0.143 | 0.143 | 0.143 |

**Supplementary Table 7 | Cryo-EM data collection and refinement statistics for 60S and 80S ribosomal states from the HEK293T cryoGO dataset**

|  | State 1a | State 1b | State 1c | State 2a | State 2b | State 2c | State 3a | State 3b | State 3c | State 4a |
| --- | --- | --- | --- | --- | --- | --- | --- | --- | --- | --- |
|  | EMD-77428 | EMD-77429 | EMD-77430 | EMD-77431 | EMD-77432 | EMD-77433 | EMD-77434 | EMD-77435 | EMD-77436 | EMD-77437 |
| **Data collection and processing** |  |  |  |  |  |  |  |  |  |  |
| Magnification | 81,000 | 81,000 | 81,000 | 81,000 | 81,000 | 81,000 | 81,000 | 81,000 | 81,000 | 81,000 |
| Voltage (kV) | 300 | 300 | 300 | 300 | 300 | 300 | 300 | 300 | 300 | 300 |
| Electron exposure (e-/ Å^2^) | 50 | 50 | 50 | 50 | 50 | 50 | 50 | 50 | 50 | 50 |
| Defocus range (μm) | -0.5 to -1.5 | -0.5 to -1.5 | -0.5 to -1.5 | -0.5 to -1.5 | -0.5 to -1.5 | -0.5 to -1.5 | -0.5 to -1.5 | -0.5 to -1.5 | -0.5 to -1.5 | -0.5 to -1.5 |
| Pixel size (Å) | 1.068 | 1.068 | 1.068 | 1.068 | 1.068 | 1.068 | 1.068 | 1.068 | 1.068 | 1.068 |
| Micrographs (no.) | 7,036 | 7,036 | 7,036 | 7,036 | 7,036 | 7,036 | 7,036 | 7,036 | 7,036 | 7,036 |
| Final particle images (no.) | 35,052 | 27,649 | 9,145 | 148,736 | 115,578 | 36,939 | 9,946 | 10,781 | 8,234 | 19,196 |
| Symmetry imposed | C1 | C1 | C1 | C1 | C1 | C1 | C1 | C1 | C1 | C1 |
| Map resolution (Å) | 2.81 | 2.88 | 3.11 | 2.38 | 2.46 | 2.73 | 3.15 | 3.11 | 3.16 | 2.96 |
| FSC threshold | 0.143 | 0.143 | 0.143 | 0.143 | 0.143 | 0.143 | 0.143 | 0.143 | 0.143 | 0.143 |

**Supplementary Table 7 continued**

|  | State 4b | State 4c | State 4d | State 5 | State 6 | State 7a | State 7b | State 8a | State 8b | State 8c |
| --- | --- | --- | --- | --- | --- | --- | --- | --- | --- | --- |
|  | EMD-77438 | EMD-77439 | EMD-77440 | EMD-77441 | EMD-77442 | EMD-77443 | EMD-77444 | EMD-77445 | EMD-77446 | EMD-77447 |
| **Data collection and processing** |  |  |  |  |  |  |  |  |  |  |
| Magnification | 81,000 | 81,000 | 81,000 | 81,000 | 81,000 | 81,000 | 81,000 | 81,000 | 81,000 | 81,000 |
| Voltage (kV) | 300 | 300 | 300 | 300 | 300 | 300 | 300 | 300 | 300 | 300 |
| Electron exposure (e-/ Å^2^) | 50 | 50 | 50 | 50 | 50 | 50 | 50 | 50 | 50 | 50 |
| Defocus range (μm) | -0.5 to -1.5 | -0.5 to -1.5 | -0.5 to -1.5 | -0.5 to -1.5 | -0.5 to -1.5 | -0.5 to -1.5 | -0.5 to -1.5 | -0.5 to -1.5 | -0.5 to -1.5 | -0.5 to -1.5 |
| Pixel size (Å) | 1.068 | 1.068 | 1.068 | 1.068 | 1.068 | 1.068 | 1.068 | 1.068 | 1.068 | 1.068 |
| Micrographs (no.) | 7,036 | 7,036 | 7,036 | 7,036 | 7,036 | 7,036 | 7,036 | 7,036 | 7,036 | 7,036 |
| Final particle images (no.) | 16,406 | 7,588 | 22,129 | 24,560 | 18,989 | 14,635 | 8,188 | 6,175 | 12,684 | 4,679 |
| Symmetry imposed | C1 | C1 | C1 | C1 | C1 | C1 | C1 | C1 | C1 | C1 |
| Map resolution (Å) | 3.00 | 3.25 | 2.89 | 2.88 | 2.98 | 2.93 | 3.40 | 3.21 | 2.99 | 3.62 |
| FSC threshold | 0.143 | 0.143 | 0.143 | 0.143 | 0.143 | 0.143 | 0.143 | 0.143 | 0.143 | 0.143 |

**Supplementary Table 7 continued**

|  | State 9a | State 9b | State 9c | State 9d | State 10a | State 10b | State 10c | State 10d | State 10e | State 10f |
| --- | --- | --- | --- | --- | --- | --- | --- | --- | --- | --- |
|  | EMD-77448 | EMD-77449 | EMD-77450 | EMD-77451 | EMD-77452 | EMD-77453 | EMD-77454 | EMD-77455 | EMD-77456 | EMD-77457 |
| **Data collection and processing** |  |  |  |  |  |  |  |  |  |  |
| Magnification | 81,000 | 81,000 | 81,000 | 81,000 | 81,000 | 81,000 | 81,000 | 81,000 | 81,000 | 81,000 |
| Voltage (kV) | 300 | 300 | 300 | 300 | 300 | 300 | 300 | 300 | 300 | 300 |
| Electron exposure (e-/ Å^2^) | 50 | 50 | 50 | 50 | 50 | 50 | 50 | 50 | 50 | 50 |
| Defocus range (μm) | -0.5 to -1.5 | -0.5 to -1.5 | -0.5 to -1.5 | -0.5 to -1.5 | -0.5 to -1.5 | -0.5 to -1.5 | -0.5 to -1.5 | -0.5 to -1.5 | -0.5 to -1.5 | -0.5 to -1.5 |
| Pixel size (Å) | 1.068 | 1.068 | 1.068 | 1.068 | 1.068 | 1.068 | 1.068 | 1.068 | 1.068 | 1.068 |
| Micrographs (no.) | 7,036 | 7,036 | 7,036 | 7,036 | 7,036 | 7,036 | 7,036 | 7,036 | 7,036 | 7,036 |
| Final particle images (no.) | 4,656 | 5,092 | 6,321 | 5,742 | 14,635 | 8,188 | 6,175 | 19,626 | 19,274 | 17,816 |
| Symmetry imposed | C1 | C1 | C1 | C1 | C1 | C1 | C1 | C1 | C1 | C1 |
| Map resolution (Å) | 3.65 | 3.55 | 3.24 | 3.34 | 2.93 | 3.40 | 3.21 | 2.82 | 2.79 | 2.81 |
| FSC threshold | 0.143 | 0.143 | 0.143 | 0.143 | 0.143 | 0.143 | 0.143 | 0.143 | 0.143 | 0.143 |
